# Live-Cell Quantification Reveals Viscoelastic Regulation of Synapsin Condensates by α-Synuclein

**DOI:** 10.1101/2024.07.28.605529

**Authors:** Huan Wang, Christian Hoffmann, Johannes V. Tromm, Xiao Su, Jordan Elliott, Han Wang, Jean Baum, Zhiping P. Pang, Dragomir Milovanovic, Zheng Shi

## Abstract

Synapsin and α-synuclein represent a growing list of condensate-forming proteins where the material states of condensates are directly linked to cellular functions (e.g., neurotransmission) and pathology (e.g., neurodegeneration). However, quantifying condensate material properties in living systems has been a significant challenge. To address this, we develop MAPAC (**<u>m</u>**icropipette **<u>a</u>**spiration and whole-cell **<u>pa</u>**tch **<u>c</u>**lamp), a platform that allows direct material quantification of condensates in live cells. We find 10,000-fold variations in the viscoelasticity of synapsin condensates, regulated by the partitioning of α-synuclein, a marker for synucleinopathies. Through in vitro reconstitutions, we identify 4 molecular factors that distinctly regulate the viscosity and interfacial tension of synapsin condensates, verifying the cellular effects of α-synuclein. Overall, our study provides unprecedented quantitative insights into the material properties of neuronal condensates and reveals a crucial role of α-synuclein in regulating condensate viscoelasticity. Furthermore, we envision MAPAC applicable to study a broad range of condensates in vivo.

## Introduction

Biomolecular condensates that form through phase separation have recently come to prominence as a key contributor to various cellular functions ^1–3^. The viscoelasticity of condensates regulates the kinetics of biochemical reactions in the dense phase, whereas an aberrant increase in condensate viscoelasticity has been associated with neurodegenerative diseases ^4–12^. Beyond bulk viscoelasticity, there has been a growing interest in the interfacial properties of condensates ^13^, focusing on their roles in regulating the structure of multiphase condensates ^14–17^, the growth and size of condensates ^18–20^, and the interactions between condensates with other cellular components such as biopolymers ^19, 21–23^ and membrane-bound organelles ^24–28^.

Neuronal communication critically relies on the regulated secretion of synaptic vesicles (SVs) at specialized sites termed synapses. The cluster of SVs at the synaptic boutons represents a prominent example of biomolecular condensates ^26, 29, 30^, assembled by the highly abundant, intrinsically disordered synapsin family of proteins ^31, 32^. Synapsin/SV condensates act as reaction centers that are able to recruit α-synuclein ^33–35^, a major synaptic protein implicated in the regulation of SV cycles and pathology of neurodegenerative diseases collectively known as synucleinopathies ^36–38^. Genetic data suggest a tight balance of synapsin and α-synuclein concentrations that directly affects the mesoscale organization of SV clusters at synapses ^39^. The altered concentration of α-synuclein disrupts SV packing ^40, 41^ and impairs neurotransmission ^34, 36, 39, 42^. Therefore, the material properties of synapsin condensates and factors that regulate these properties can directly affect neuronal signaling and neurodegenerative diseases.

While previous data provide circumstantial evidence that α-synuclein plays a role in the organization of synapsin/SV assemblies, direct measurements are lacking to determine whether and how α-synuclein regulates the material state of synapsin condensates due to technical limitations. Recently, we demonstrated that micropipette aspiration (MPA) can be applied to quantify the material properties of condensates ^43, 44^. MPA is a label-free technique that directly measures both the viscosity and interfacial tension of condensates ^43, 45^. This differentiates MPA from widely used techniques such as fluorescence recovery after photobleaching (FRAP), fluorescence correlation spectroscopy (FCS), microrheology, and droplet fusion, where either only the viscosity or a ratio of viscosity to interfacial tension of condensates can be probed ^46^. However, it remains challenging to quantify condensate material properties in living systems ^12, 47, 48^, thus severely hindering our understanding of condensates in their native environment.

Here, we develop a correlative **M**P**A** and whole-cell **pa**tch **c**lamp (MAPAC) platform to study the material properties of synapsin condensates in living mammalian cells. The recording of cell membrane voltage provides an electrophysiological monitor for cell health ^49^ and aids in the detection of condensates in the cytosol. Interestingly, the viscous and elastic moduli of condensates vary by 4 orders of magnitude between cells, and these variations are directly correlated with the partitioning of α-synuclein, and, to a lesser extent, the partitioning of synapsin into condensates. Furthermore, we explore the material properties of reconstituted synapsin condensates in vitro. We show that removing the N-terminal region of synapsin leads to a reduction in condensate viscosity. In contrast, α-synuclein and two additional factors (crowding agents and SVs) significantly increase the viscosity of synapsin condensates. However, these factors have distinct effects on the interfacial tension of synapsin condensates: the interfacial tension increases with PEG, decreases with SVs, but remains unaltered with the partitioning of α-synuclein. The in vitro results directly verify our observed effect of α-synuclein on the viscosity and interfacial tension of synapsin condensates in live cells. In contrast to the viscoelastic condensates observed in cells, reconstituted synapsin condensates appear purely viscous, suggesting additional cellular factors that promote condensate elasticity in a living system. Together, our results provide unprecedented quantitative understanding of synapsin condensates, highlighting an effective cellular control of condensate viscoelasticity via compositional modulations. Moreover, we present MAPAC, a broadly applicable platform that allows intricate examinations of individual condensates in living cells.

## Results

### MAPAC allows direct measurements of synapsin condensates in live cells

To probe condensates in live cells, an electrode-containing micropipette filled with intracellular buffer solution was positioned to break the plasma membrane near a condensate (Figure 1A). Once the locally broken plasma membrane sealed to the edge of the pipette tip, suction pressures were gently applied so that the nearby condensate could flow into the tip of the micropipette for subsequent quantifications. Meanwhile, the recorded membrane voltage served as a sensitive monitor for the stability of the intracellular environment during MAPAC.

**Figure 1:**
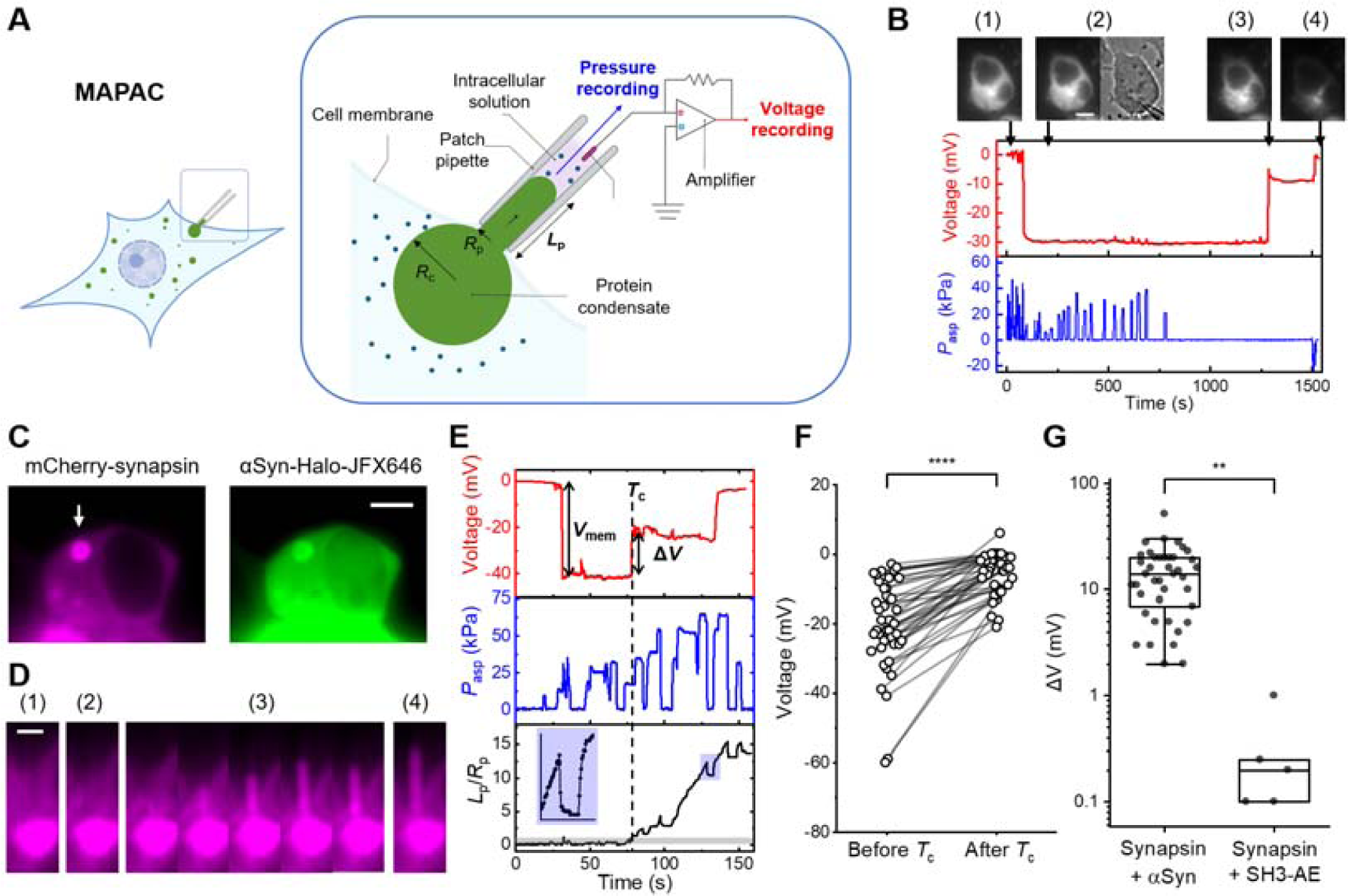
Development of MAPAC for condensate measurements in live cells. **(A)** Schematic representation of the MAPAC platform. The subcellular region where the micropipette broke into the cell and aspirated a condensate is zoomed-in on the right. **(B)** An example testing the effect of aspiration pressure (blue) on cell membrane voltage recording (red) on a HEK 293T cell expressing mCherry-synapsin and α-synuclein-BFP. Four fluorescence images of synapsin correspond to timepoints before (1) and after (2) break-in; before (3) and after (4) losing membrane seal. A transmitted light image is included for timepoint (2) to show the patched cell and the position of the micropipette. The voltage signal before timepoint (3) corresponded to a motion of the micropipette tip that either contacted the condensate or weakened the membrane seal. The voltage jumped to 0 mV before timepoint (4) due to an intentional ejection pressure that broke the membrane seal. Significant leakage of cytosolic fluorescence was observed after the ejection pressure. **(C)** Fluorescence images showing a HEK 293T cell co-expressing mCherry-synapsin (magenta) and α-synuclein-Halo (stained with JFX-646; green). The arrow represents the micropipette that was positioned next to a condensate. **(D)** Fluorescence images of the synapsin/α-synuclein condensate before (1) and after (2) micropipette break-in, during MAPAC (3), and after *V*_mem_ jumped to zero (4). **(E)** MAPAC recording of membrane voltage (red), aspiration pressure (blue), and aspiration length (black; normalized to pipette radius) for the condensate in **(D)**. The entrance of the condensate into pipette tip at *T*_c_ (dashed line) accompanied a voltage jump (Δ*V*). The grey shade on *L*_p_/*R*_p_ represents *L*_p_/*R*_p_ < 1. The blue shade on *L*_p_/*R*_p_ is zoomed-in on the left, showing viscoelastic responses of the condensate to pressure steps. **(F)** Voltage recordings before and after a synapsin/α-synuclein condensate entered the tip of the micropipette. *n* = 41 cells. **** *p* < 10^−4^, paired *t* test. **(G)** Δ*V* measured on synapsin/α-synuclein condensates and synapsin/intersectin-SH3 condensates (n = 5). ** *p* < 10^−2^, Student’s *t* test. All scale bars are 5 μm, except 2 μm in **(D)**.

First, we tested the effect of aspiration pressure (*P*_asp_) on the stability of voltage recording. Under frequent pressure perturbations that resemble those used to probe condensate material properties, the measured membrane voltage on human embryonic kidney (HEK) 293T cells was stable for tens of minutes until purposefully perturbed by an ejection pressure that broke the membrane seal (Figure S1A). A similar time window was maintained in cells over-expressing synapsin and α-synuclein (Figure 1B), sufficient for extensive aspiration measurements where each pressure step takes ∼ 10 seconds. In addition to membrane voltage, the fluorescence of cytosolic proteins can be used as a secondary monitor for potential leakages at the cell membrane (Figure 1B).

In HEK 293T cells, the expression of synapsin alone was not sufficient to drive condensate formation, likely due to the lack of client proteins and other neuronal components naturally occurring to promote phase separation at the nerve terminal ^50^. Indeed, co-expressing synapsin with α-synuclein or intersectin SH3 domains robustly induced micrometer-size synapsin condensates (Figure 1C, Figure S1) ^33^. We focused our further study on a minimal system with HEK 293T cells co-expressing fluorescently tagged full-length synapsin 1 and α-synuclein due to their relevance in neurotransmission and synucleinopathies ^33^ (Figure 1C). These cells typically have one (occasionally two) major condensate with a radius *R*_c_ = 1.5 ± 0.6 μm (mean ± SD, n = 74; Figure S1B), sufficient for accurate aspiration measurements with patch-clamp micropipettes that have an opening diameter ∼1 μm (Figure 1D, Movie S1).

After breaking into the cell but before aspirating a condensate, the electrode-containing micropipette measures the resting voltage of the cell membrane (*V*_mem_; Figure 1E). On non-transfected cells, the recorded *V*_mem_ = −49 ± 6 mV (mean ± SEM, n = 7; Figure S1C) was consistent with membrane potentials reported for healthy HEK 293T ^51^. Cells expressing synapsin and α-synuclein showed a less polarized membrane potential (*V*_mem_ = −23 ± 2 mV; mean ± SEM, n = 47; *p* < 0.01; Figure S1C), indicating an electrophysiological perturbation by the expressed neuronal proteins. Interestingly, a voltage jump (Δ*V*) was observed when synapsin/α-synuclein condensates entered the tip of the micropipette (Figure 1E). The measured Δ*V* = 15 ± 1.5 mV (mean ± SEM, n = 41; *p* < 10^−4^; Figure 1F) might suggest that synapsin/α-synuclein condensates significantly increased the resistance to ion flow into the micropipette ^52–54^. Δ*V* was more significant for synapsin condensates driven by α-synuclein than those driven by intersectin SH3 domains (Figure 1G; *p* < 0.01), indicating different porosity ^55^, thus electrical resistance, of these condensates. On a technical note, Δ*V* can be used to aid label-free detections of condensates by the micropipette tip.

### Synapsin condensates in cells are viscoelastic

After a condensate entered the tip of the micropipette, controlled steps of aspiration pressure (*P*_asp_) were applied to examine the material properties of the condensate (Figure 1E; Movie S1). Most cellular synapsin condensates showed clear features of viscoelasticity, as indicated by sudden nonlinear jumps of aspiration length (*L*_p_) upon stepwise pressure changes (Figure 1E, Figure S1D – S1G). Additionally, cellular condensates did not wet the inner wall of micropipettes: condensates inside the micropipette formed a convex meniscus, could undergo necking instability, and did not leave behind any measurable fluorescence after leaving the micropipette (Figure S2). Consistent with a slipping contact between the condensate and the micropipette, the long-term response of *L*_p_ under constant pressure appeared linear (Figure 1E, 2A)^44, 56^.

**Figure 2:**
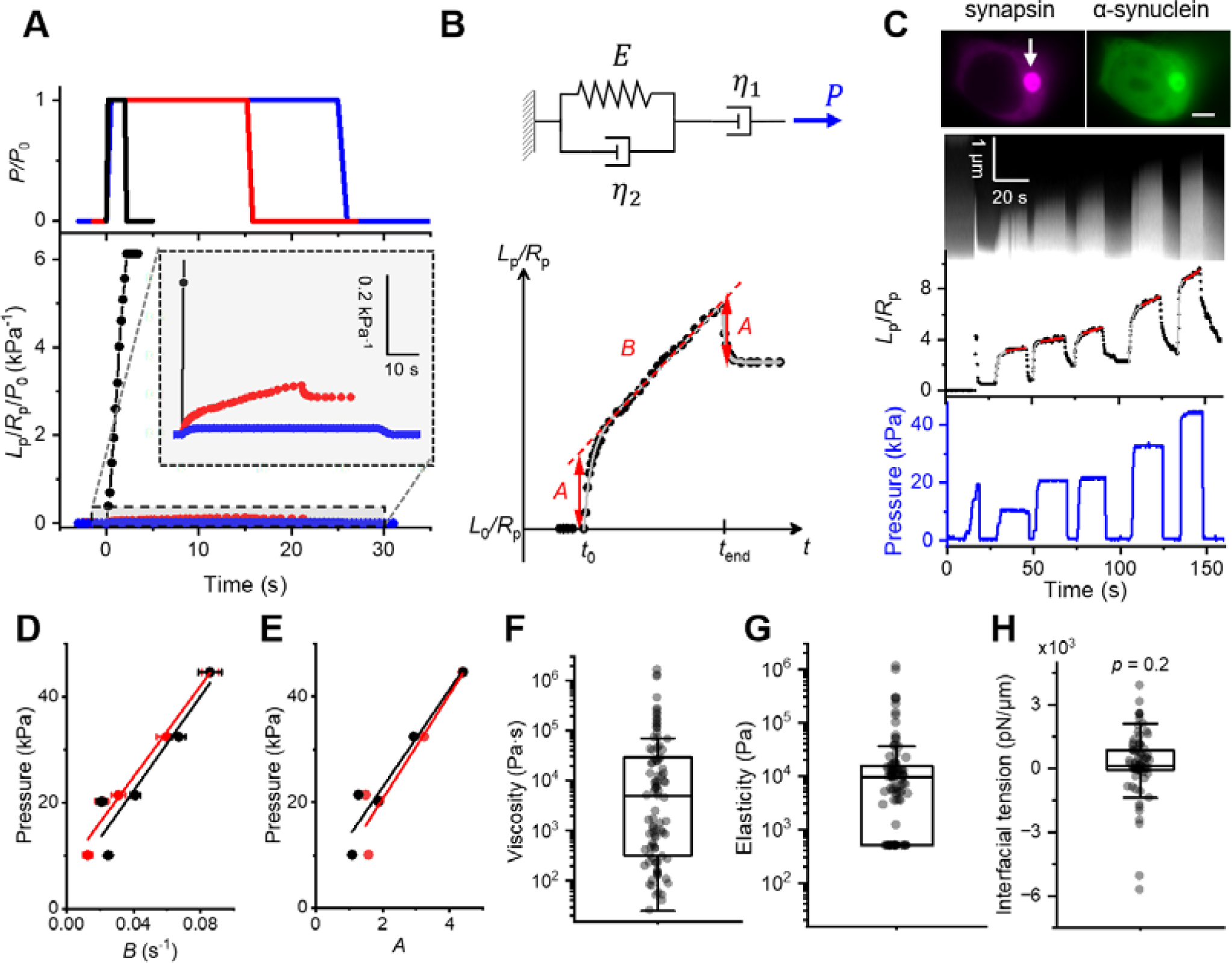
Cellular synapsin condensates quantified by a 3-element viscoelastic liquid model. **(A)** Representative responses of cellular synapsin condensates to a stepwise aspiration pressure. Black: apparent viscous liquid, 24/93 condensates; red: viscoelastic liquid, 53/93 condensates; blue: apparent elastic solid, 16/93 condensates. The boxed area is zoomed-in to improve the display of the red and blue data. **(B)** Upper: the 3-element viscoelastic liquid model used for analyzing all cellular condensates. Lower: fitting of a representative condensate response to the full model (3-parameter; grey) or to a simplified linear model (2-parameter, red). *B* is the long-term shear rate; *A* is the normalized amplitude of the elastic response. **(C)** Upper: fluorescence images of a HEK 293T cell co-expressing mCherry-synapsin (magenta) and α-synuclein-Halo labeled with JFX-646 (green), arrow points to the condensate, scale bar 5 μm. Middle: kymograph of the aspirated part of the condensates. Lower: normalized aspiration length (black), in response to stepwise aspiration pressure (blue). Grey curves are fits to the full model (eq. 1); red lines are linear fits. **(D-E)** Aspiration pressures (*P*_asp_) plotted against either the long-term shear rate *B* (**D)** or the normalized amplitude of elastic response *A* (**E)**. Black and red markers represent results from the full model and the linear model, respectively. Lines are linear fits. **(F-H)** Material properties of synapsin condensates in cells. Viscosity **(F)** was calculated from the slope of **D** (n = 93); elasticity **(G)** was calculated from the slope of **E** (n = 93), 24 condensates with apparent pure viscous response were assigned an elasticity of 500 Pa that corresponded to the experimental resolution; interfacial tension **(H)** was calculated from the intercepts of **D** and **E** (n = 77). *p* value from one sample t-test.

Across ∼ 100 synapsin/α-synuclein condensates in HEK 293T cells, we observed material responses ranging from apparent viscous liquid (Figure 2A, black) to viscoelastic liquid (Figure 2A, red) and to apparent elastic solid (Figure 2A, blue). We found that a 3-element viscoelastic liquid (Jeffreys) model is minimally required to explain all condensate responses observed in cells (Supplementary Discussions) ^57^. The model includes a viscous liquid component (η_1_) that governs the long-term condensate response, in series with a viscoelastic solid component (*E* and η_2_) that determines the short-term response (Figure 2B) ^56^. Under constant pressure, the time-dependent *L*_p_ (normalized to the micropipette radius *R*_p_) follows equation (1):

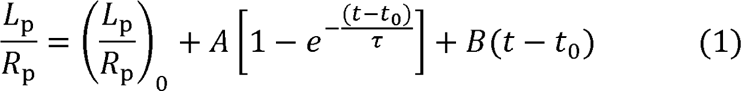

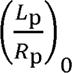 is the normalized aspiration length before a stepwise pressure change at time *t*_0_. The model has three fitting parameters: 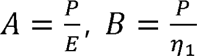, and 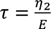. Here, *P* = *P*_asp_ - *P*_γ_, and 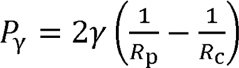 s the capillary pressure that needs to be overcome to initiate condensate flow. In this model, an apparent elastic condensate (Figure 2A, blue) will have an infinite viscosity, whereas an apparent viscous condensate (Figure 2A, black) can have zero elasticity. We assigned these extreme cases using the resolution of our experiments (Supplementary Discussions).

Next, we measured the “*L*_p_/*R*_p_” of condensates under systematically varied steps of *P*_asp_ (Figure 2C; Movie S2). The slopes of “*P*_asp_ vs. *B*” and “*P*_asp_ vs. *A*” give η_1_ and *E*, respectively, whereas the intercepts give *P*_γ_ (Figure 2D, 2E). η_1_, *E*, and *P*_γ_ can then be converted to the viscosity, elasticity, and interfacial tension of the measured condensate, respectively (Supplementary Discussions) ^56, 58^. Notably, a 2-parameter linear model can be applied to the long-term response of *L*_p_ (Figure 2B, 2C) ^56^. The resulting “*P*_asp_ vs. *B*” and “*P*_asp_ vs. *A*” relations were indistinguishable from those of the full 3-parameter model (Figure 2D, 2E, Figure S3). By sacrificing the fitting for η_2_ (Figure S3F), a property not directly related to the material state of the condensate ^57^, the linear model reduced fitting errors, especially for data with low signal to noise ratio and for data where the pressure profile significantly deviated from a step-function (Figure S3). Therefore, we used the linear model for all further analysis of cellular condensates.

Cellular synapsin condensates exhibited viscoelastic moduli that span over 4 orders of magnitude (Figure 2F, 2G; n = 93) from liquid states (viscosity < 10^2^ Pa·s) to apparent solid states that cannot be measurably deformed (elasticity ∼ 10^6^ Pa). The viscoelasticity of cellular condensates were intrinsic properties of the condensate-forming proteins, as they did not depend on the resting potential of the cell and were not significantly affected by the choice of fluorescent tags (Figure S4). The measured interfacial tension of cellular condensates (γ = 240 ± 220 pN/μm; mean ± SEM; Figure 2H, Figure S5) was not significantly different from zero (*p* = 0.2), consistent with the low interfacial tension of protein condensates in vitro ^43^. The uncertainties in interfacial tension correspond to a capillary pressure *P*_γ_ (∼ 1 kPa) that did not significantly affect aspiration pressures used for viscoelasticity measurements (10 ∼ 50 kPa; Figure 2C).

The viscosity of cellular synapsin condensates (median = 5000 Pa·s; Figure 2F) was significantly higher than the viscosity of protein condensates measured in vitro (1 – 1000 Pa·s) ^43, 59^. However, cellular condensates that appear highly viscoelastic may resemble disease-related solid aggregates such as α-synuclein containing Lewy bodies ^60, 61^. Therefore, we next explore whether the partitioning of α-synuclein plays a role in determining the viscoelasticity of synapsin condensates in cells.

### Condensate partitioning of α-synuclein dictates the viscoelasticity of synapsin condensates in cells

When α-synuclein and synapsin were heterogeneously expressed in HEK 293T cells, the condensate partitioning coefficient of α-synuclein (π_αSyn_; i.e., the concentration of When α-synuclein and synapsin were heterogeneously expressed in HEK 293T cells, magnitude, from weakly depleted from condensates (π_αSyn_ < 1) to highly enriched in α-synuclein in condensates relative to that in the dilute phase) spanned two orders of the condensate phase (π_αSyn_ > 10; Figure S6A – S6C). In comparison, the condensate partitioning coefficient of synapsin (π_synapsin_) was more clustered (∼ 10) and overall higher than π_αSyn_ (*p* < 10^−4^; Figure S6C), reminiscent to the observation of α-synuclein being a client protein recruited to condensates formed by synapsin ^33^. For synapsin/α-synuclein condensates with comparable π_synapsin_ (10 ± 3), we found π_αSyn_ to be an accurate prophecy of the condensate’s viscoelastic response under ∼ 5 condensates was ∼100 times slower; whereas condensates with π_αSyn_ > 10 were barely deformable (Figure 3A). Across 43 cells that co-expressed mCherry-synapsin condensates strongly correlated with π_αSyn_ (Figure 3B, Figure S6D; Pearson’s r = and α-synuclein-Halo (stained with JFX-646), the viscoelasticity of formed 0.84 for viscosity, r = 0.74 for elasticity). In contrast, the viscoelasticity was not dependent on the apparent concentrations of α-synuclein in the cytosol (r = 0.08; Figure S6E) and was only weakly dependent on the apparent concentration of α-synuclein in the condensate (r = 0.39; Figure S6F), consistent with the partitioning coefficient being a better indicator of α-synuclein’s affinity towards the condensates (Supplementary Discussions) ^12^. Additionally, the measured viscoelasticity and partitioning coefficients were independent of condensate radius (r = 0.14; Figure S6G), confirming the lack of optical artifacts induced by condensate curvature.

**Figure 3:**
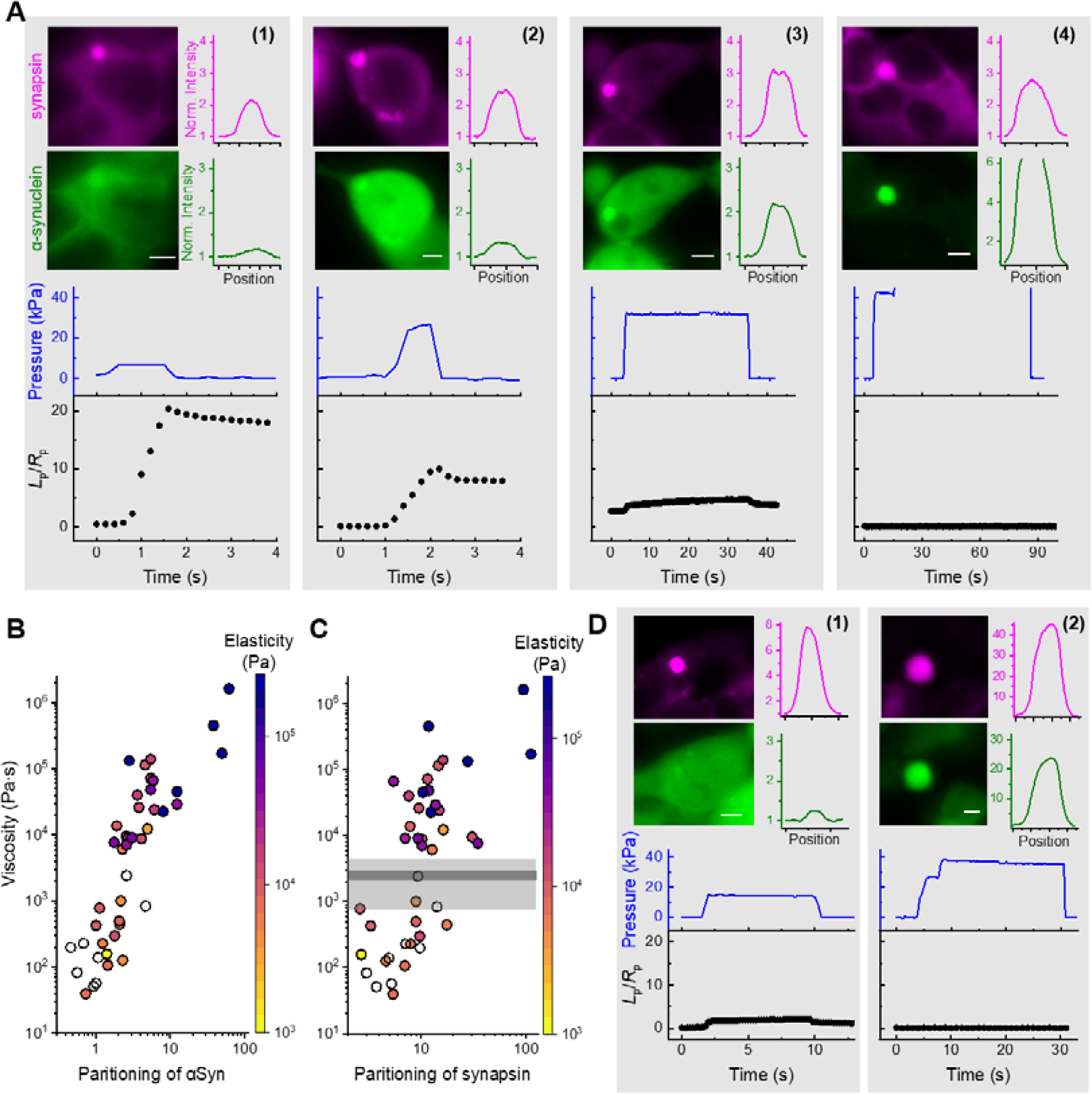
The partitioning of α-synuclein dictates the viscoelasticity of synapsin condensate in cells. **(A)** Upper: widefield fluorescence images of four HEK 293T cells co-expressing mCherry-synapsin (magenta) and α-synuclein-Halo labelled with JFX-646 (green). Fluorescence intensity profiles across the condensate are plotted on the right of each image. These four condensates have similar partitioning of synapsin but drastically different partitioning of α-synuclein, from (1) to (4): *π*_synapsin_ = 5.2, 9.7, 13.7, 11.7; and *π*_αSyn_ = 1.0, 1.8, 5.5, 38, respectively. See Figure S6 for conversion from apparent partitioning measured from widefield to true partitioning measured from confocal microscopy. Lower: applied aspiration pressure step (blue) and corresponding response of each condensate (black). All graphs have the same scale in *P*_asp_ and *L*_p_/*R*_p_. The condensate increased in viscosity and elasticity from (1) to (4). **(B, C)** The relationships between the viscosity of 43 condensates as a function of *π*_αSyn_ **(B)** or *π*_synapsin_ **(C)**. The elasticity of each condensate was colored according to the map. White condensates did not show measurable elastic response and was assigned an elasticity of 500 Pa. The gray line and shade represent the mean and standard deviation of the system viscosity (*η*_2_/6) measured from the full model. **(D)** Upper: widefield fluorescence images of two HEK 293T cells co-expressing mCherry-synapsin (magenta) and α-synuclein-Halo labelled with JFX-646 (green), with fluorescence intensity profiles across the condensate plotted on the right of each image. From (1) to (2): *π*_synapsin_ = 35, 93; and *π*_αSyn_ = 1.8, 61, respectively. Lower: applied aspiration pressure step (blue) and corresponding response of each condensate (black). Error bars are SEM. All scale bars are 5 μm.

A positive correlation (r = 0.68) was observed between π_αSyn_ and π_synapsin_ in cells cooperativity between α-synuclein and synapsin. Consequently, π_synapsin_ can also (Figure S6H), indicating the presence of cellular factors that promoted heterotropic 0.66 for viscosity, r = 0.60 for elasticity) albeit less accurately compared to π_αSyn_. predict the viscoelasticity of cellular condensates (Figure 3C, Figure S6I – S6K; r = Condensates with π_αSyn_ ∼ 1 but π_synapsin_ > 30 showed strong viscoelastic features, while condensates with high values (> 30) of both π_αSyn_ and π_synapsin_ were not measurably deformable (Figure 3D). Linear combinations of π_αSyn_ and π_synapsin_ did not outperform π_αSyn_ alone in predicting the viscoelasticity of cellular condensates (Figure S6L), consistent with the partition of α-synuclein being the main driver for the viscoelasticity of synapsin condensates in cells.

### Synapsin domains and crowding conditions regulate the viscosity and interfacial tension of synapsin condensates

To verify the effect of α-synuclein and to explore additional regulators on the material properties of synapsin condensates, next we explore synapsin condensates reconstituted in vitro ^26^. Full-length synapsin 1 (hereafter named “FL synapsin”) comprises a membrane binding domain and an ATP-binding domain at the N-terminal, with an intrinsically disordered region (IDR) at the C-terminal (Figure 4A) ^26, 31^. Purified FL synapsin spontaneously forms condensates under physiologically relevant conditions: 5 ∼ 10 μM synapsin in 150 mM NaCl, 3 ∼ 10% polyethylene glycol-8000 (hereafter named “PEG”), pH 7.4. While our cellular measurements (Figure 1 – 3) focused on FL synapsin, in vitro studies have identified the IDR to be the main driver of the phase separation of synapsin: deleting IDR abolishes phase separation, while the purified IDR fragment can form condensates on its own ^26^. However, it is unclear whether condensates formed by the IDR are distinct, in their material properties, from those consisting of FL synapsin.

**Figure 4:**
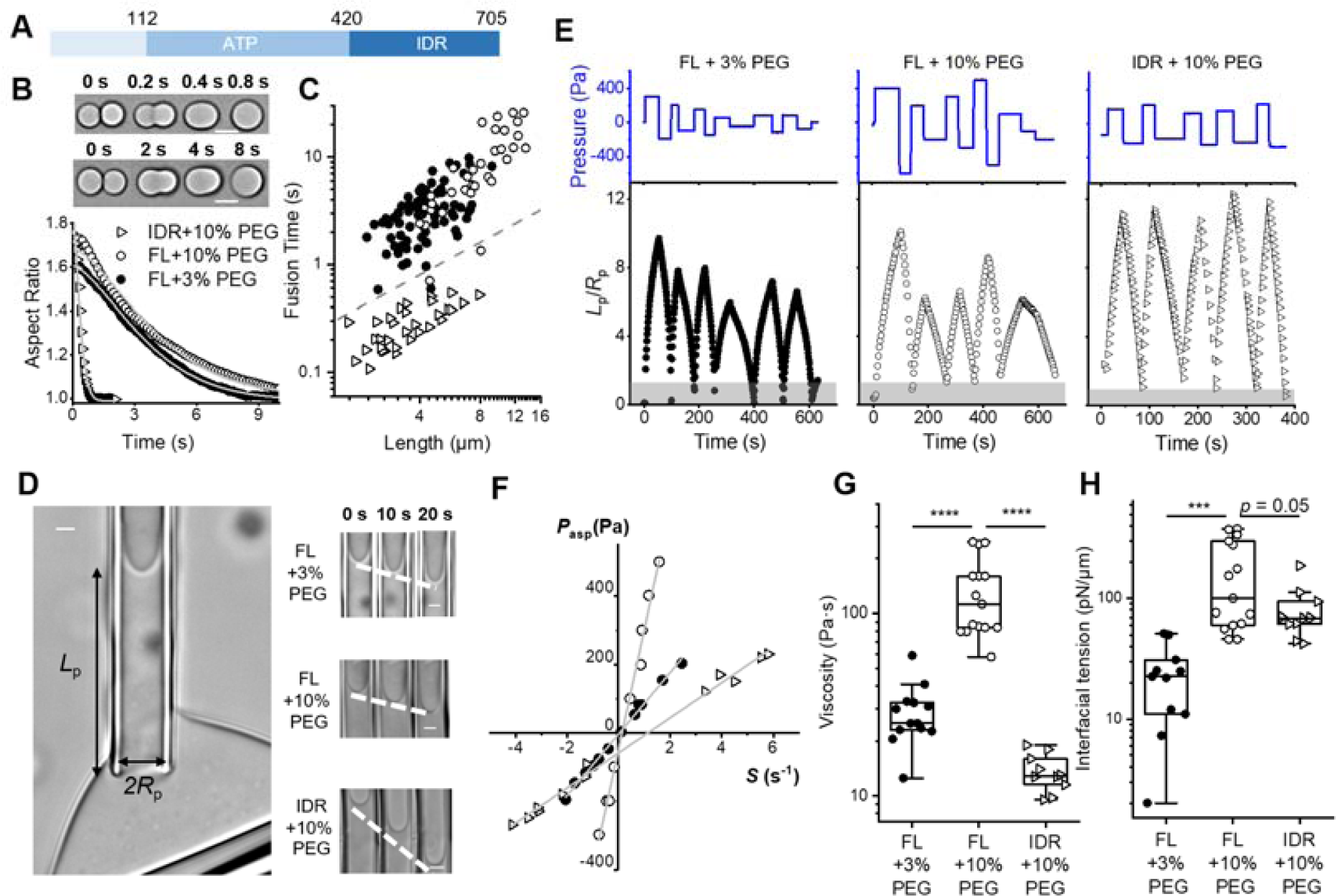
PEG increases both the viscosity and interfacial tension of synapsin condensates while the IDR of synapsin maintains a low condensate viscosity. **(A)** Domain diagram of full-length (FL) synapsin 1. **(B)** Images showing the fusion of IDR (upper) and FL (middle) synapsin condensates. Lower: the relaxation of condensate aspect ratios. Grey curves are stretched-exponential fits. Open triangles: IDR under 10% PEG; open circles: FL under 10% PEG; close circles: FL under 3% PEG; same below. Scale bars, 5 μm. **(C)** Condensate fusion time vs. condensate length (the geometric mean of condensate diameters before fusion). The inverse of the slopes from linear fitting gave capillary velocities: 13.0 ± 0.6 μm/s (IDR + 10% PEG; R^2^ = 0.93; n = 33 pairs), 0.8 ± 0.1 μm/s (FL + 10% PEG; R^2^ = 0.81; n = 34 pairs), and 1.3 ± 0.1 μm/s (FL + 3% PEG; R^2^ = 0.82; n = 76 pairs). The dashed line has a slope of 1 in log-log scale, demonstrating the expected slope for a Newtonian liquid. **(D)** Left: transmitted light image showing a micropipette-aspirated synapsin condensate (held by a second pipette). Right: time-lapse images showing the flow of FL (upper: + 3% PEG; middle: + 10% PEG) and IDR (lower: + 10% PEG) synapsin condensates under constant aspiration pressure (upper and middle: *P*_asp_ = −300 Pa; lower: *P*_asp_ = −250 Pa). *P*_asp_ < 0 corresponds to ejection. All condensates wet the inner wall of micropipettes as indicated by the concave meniscus. Scale bars, 2 μm. **(E)** Aspiration pressure (blue) and normalized aspiration length (black) during MPA of FL (left: +3% PEG; middle: +10% PEG) and IDR (right: +10% PEG) synapsin condensates. The grey shades represent *L*_p_/*R*_p_ < 1. **(F)** The relationship between aspiration pressure and shear rate *S*, defined as *S* = d(*L*_p_/*R*_p_)^2^/d*t*, for synapsin condensates. Solid lines are linear fits (all R^2^ > 0.99). **(G)** Viscosity and **(H)** interfacial tension of FL (+ 3% PEG, n = 13; + 10% PEG, n = 15) and IDR (+ 10% PEG, n= 10) synapsin condensates measured by MPA. All measurements were done with 9 μM synapsin (FL or IDR) in 150 mM NaCl, 25 mM Tris-HCl, 0.5 mM TCEP, pH 7.4. *p* values were determined using one-way ANOVA followed by *post hoc* Tukey’s test. \*\*\*\**p* < 10^−4^; \*\**p* < 0.01. In the box plots, the central lines denote the median, the boxes span the interquartile range (25%-75%), and the whiskers extend to 1.5 times the interquartile range, same below.

Using an optical tweezers-assisted droplet fusion assay (Figure 4B) ^43, 46^, we found the capillary velocity of IDR condensates (13.0 ± 0.6 μm/s) was more than 10 times faster than that of the FL condensates (0.8 ± 0.1 μm/s; Figure 4C). Notably, reducing the amount of crowding agent (from 10% to 3% PEG) minimally affected the capillary velocity of synapsin condensates (Figure 4B, 4C).

The capillary velocity, which reports a ratio of interfacial tension to viscosity of the studied condensates, does not provide a direct measure of either material property ^46^. To address this, we applied MPA to synapsin condensates. All reconstituted synapsin condensates wetted the inner wall of micropipettes (Figure 4D) and the aspiration length (*L*_p_) changed with the square root of time (Figure S7). Consistent with the droplet fusion assay (Figure 4C), the responses of reconstituted synapsin condensates were not significantly different from those predicted for purely viscous liquids ^44^, suggesting condensate elasticities below 10 Pa (Supplementary Discussions). By analyzing synapsin condensates under various aspiration pressures (*P*_asp_; Figure 4E), we observed linear relations between the aspiration pressure and the effective shear rate (*S*; Figure 4F). The slope of *P*_asp_ vs. *S* reports condensate viscosity (η; Figure 4G) while the intercept reports the interfacial tension of condensates (γ; Figure 4H).

Under the same crowding condition (10% PEG), the viscosity of IDR condensates (η = 12.9 Pa·s; median, same below) was about 10 times lower than that of FL condensates (112 Pa·s; *p* <10^−4^), whereas the interfacial tensions (γ) of the two types of condensates were comparable (*p* = 0.05). These results are consistent with the ∼10-fold faster capillary velocity (γ/η) of IDR than that of FL synapsin condensates (Figure 4C). However, with direct measurements by MPA, we now assign a clear role of the synapsin N-terminal region as a mediator of the condensate’s viscosity, rather than their interfacial tension. The ∼10-fold difference between the viscosity of FL and IDR synapsin condensates is significantly larger than expectations based on the proteins’ molecular mass (FL: 102.4 kDa, IDR: 57.6 kDa; both tagged with eGFP) ^62^, consistent with the N-terminal region’s ability to dimerize synapsin ^63^.

PEG is a commonly used crowding agent that facilitates phase separation of proteins ^64, 65^. We observed that reducing the concentration of PEG from 10% to 3% led to significant decreases in both the viscosity (η from 110 to 25 Pa·s; *p* < 10^−4^, Figure 4G) and interfacial tension (γ from 100 to 23 pN/μm; *p* < 10^−3^, Figure 4H) of synapsin condensates. However, the capillary velocity (γ/η = 0.89 μm/s in 10% PEG, γ/η = 0.91 μm/s in 3% PEG) of condensates remained largely unaltered. Therefore, only measuring the capillary velocity (i.e., via droplet fusion) would have missed the significant effects of PEG on condensate material properties (for comparison, see Figure 4C). Notably, PEG has been shown to reduce the fluidity of nucleoprotein condensates ^64, 65^, suggesting a universal crowder-mediated intermolecular attraction in protein condensates ^64–67^.

### α-Synuclein increases the viscosity of synapsin condensates in a self-catalytic manner

To verify the observed effect of α-synuclein on the viscoelasticity of synapsin condensates in cells (Figure 3), we added purified α-synuclein to FL synapsin condensates in vitro. As previously reported, α-synuclein colocalized with synapsin condensates (Figure 5A) ^33^. The partitioning of α-synuclein did not affect the wetting of condensates to micropipettes (Figure 5B), and synapsin/α-synuclein condensates remained purely viscous within the timescale of our MPA measurements (Figure 5C, Figure S8A, S8B). However, the viscosity of condensates significantly increased in the presence of α-synuclein (*p* < 10^−3^; Figure 5D), whereas the interfacial tension of synapsin condensates was not significantly perturbed (*p* = 0.06; Figure 5E). The effects of α-synuclein on the viscosity and interfacial tension of reconstituted synapsin condensates directly verify our observations in cells (Figure 3B, Figure S5). The increase in condensate viscosity suggests that α-synuclein reduces the dynamics of intermolecular contact within the condensate phase ^7^, consistent with the reported attenuation of SV recycling in neurons that overexpress α-synuclein ^34, 42, 68^. Notably, this effect was nonlinear: 3 μM α-synuclein caused minimal viscosity increase (*p* = 0.13), while a ∼50-fold increase in condensate viscosity was observed in the presence of 9 μM α-synuclein (Figure 5D).

**Figure 5:**
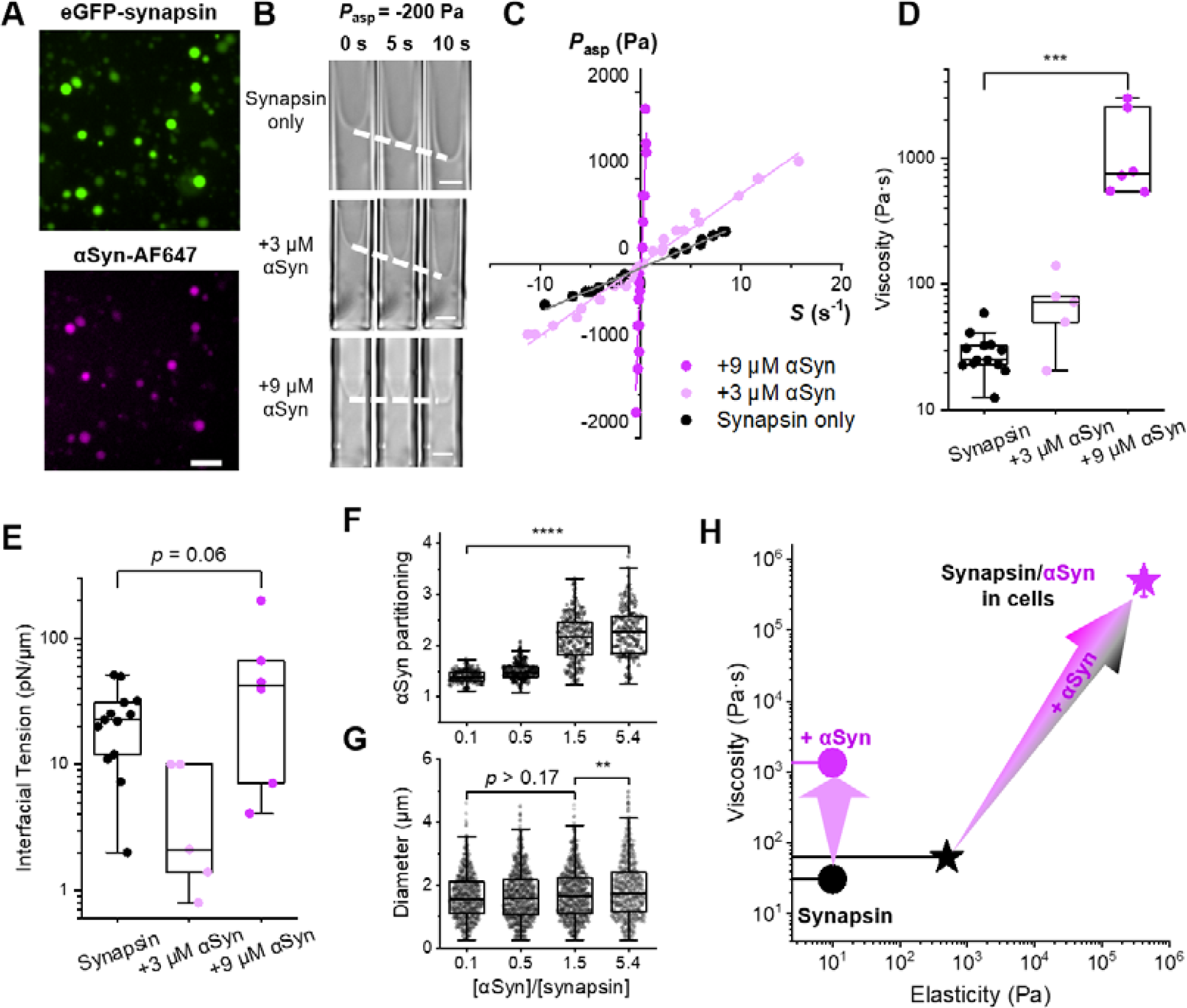
α-Synuclein increases the viscosity but does not significantly perturb the interfacial tension of full length synapsin condensates. **(A)** Colocalization of eGFP-synapsin (green) and α-synuclein (magenta; wild type α-synuclein mixed with Alexa Fluor 647 labelled α-synuclein-A140C in a 10:1 ratio). Scale bar: 10 μm. **(B)** Time-lapse transmitted light images of synapsin condensate with increasing concentration of α-synuclein. Dashed lines trace the change in *L*_p_ under −200 Pa aspiration pressure. Scale bars: 2 μm. **(C)** Relation between aspiration pressure and shear rate measured on condensates of synapsin only (black), synapsin + 3 μM α-synuclein (light magenta), and synapsin + 9 μM α-synuclein (magenta). The solid lines are linear fits. Effect of α-synuclein on the viscosity **(D)** and interfacial tension **(E)** of synapsin condensates. **(F)** The partitioning of α-synuclein to synapsin condensates with increasing molar ratio of α-synuclein to synapsin. **(G)** The diameter of synapsin condensates with increasing molar ratio of α-synuclein to synapsin. **(H)** Viscosity and elasticity of synapsin condensates. Circles represent in vitro condensates (black: synapsin only; magenta: synapsin + 9 μM α-synuclein) and were assigned an elasticity of 10 Pa. Stars represent synapsin/α-synuclein condensates in cells with low (black) and high (magenta) partitioning of α-synuclein. All synapsin proteins used in the current figure are full length, with a concentration of 9 μM in **(A)−(E)** and 5 μM in **(F)**, **(G).** Buffer: 150 mM NaCl, 25 mM Tris-HCl, 0.5 mM TCEP, pH 7.4. *p* values were determined using one-way ANOVA followed by *post hoc* Tukey’s test. \*\**p* < 0.01; \*\*\**p* < 10^−3^; \*\*\*\**p* < 10^−4^.

Under low concentrations of α-synuclein, the protein’s affinity towards synapsin condensates can be evaluated via its condensate partitioning coefficient (π_αSyn_; Supplementary Discussions) ^12^. π_αSyn_, albeit larger than 1, was much lower than the condensate partitioning coefficient of synapsin (π_synapsin_) (Figure 5F, Figure S9), consistent with the partitioning of these two proteins in cells (Figure S6C). Notably, π_αSyn_ increased with α-synuclein concentration (Figure 5F), suggesting a homotropic condensates (Supplementary Discussions). Meanwhile, π_synapsin_ remained constant cooperativity between α-synuclein molecules in their partitioning to synapsin with increasing amount of α-synuclein (Figure S9D). This cooperativity between α-synuclein is in line with the protein’s nonlinear effect on the viscosity of synapsin condensates (Figure 5D). The size distribution of condensates represents a balance between condensate interfacial tension and the mixing entropy of the system ^17^. The size of synapsin/α-synuclein condensates remained constant under a wide range of α-synuclein concentrations (Figure 5G), in agreement with the lack of α-synuclein effect on the interfacial tension of synapsin condensates in vitro (Figure 5E) and in cells (Figure S5).

While the presence of α-synuclein significantly increased the viscosity of synapsin condensates both in vitro and in live cells, cellular condensates exhibited unique elastic properties (Figure 5H). The upper limit for the elasticity of in vitro synapsin condensate (10 Pa) was 3 orders of magnitude lower than the median elasticity of synapsin condensates in cells (10 kPa), directly demonstrating the importance of live cell measurements in understanding the material properties of condensates in their natural environment.

### Synaptic vesicles increase the viscosity but lower the interfacial tension of synapsin condensates

The main physiological role of synapsin condensates is to organize the reserve pool of SVs in the presynaptic terminal ^26^. Therefore, the material state of synapsin condensates may directly impact SV recycling and neurotransmitter release that are crucial for synaptic transmission. We found that SVs purified from rat brains (visualized with the lipophilic FM 4-64 dye) partition strongly into synapsin condensates (Figure 6A) ^29^. Physiological concentrations of SVs (∼ 50 nM vesicles ^33^) significantly increased the viscosity (*p* = 0.01) but decreased the interfacial tension (*p* < 0.01) of synapsin condensates (Figure 6B – 6E, Figure S8C, S8D). The increase in condensate viscosity is consistent with SV’s ability to provide multivalent interactions that promote the phase separation of synapsin. With 46 nM SVs, some synapsin/SV condensates became irregularly shaped (Figure S8B), indicating a transition into viscoelastic states for those condensates. The increase of condensate viscosity by SVs suggests that the partitioning of SVs may slow down the transport of vesicles in the reserve pool, consistent with recent single molecular tracking measurements ^29^. The reduction in the interfacial tension agrees with SVs’ tendency to enrich to the interface of synapsin condensates ^29^, indicating a role of SVs in mediating the morphological integrity of the reserve pool. Notably, measurements on condensates formed by a ternary mixture of synapsin/α-synuclein/SV suggested weak competitions between the effects of α-synuclein and SV on synapsin condensates (Figure S10).

**Figure 6:**
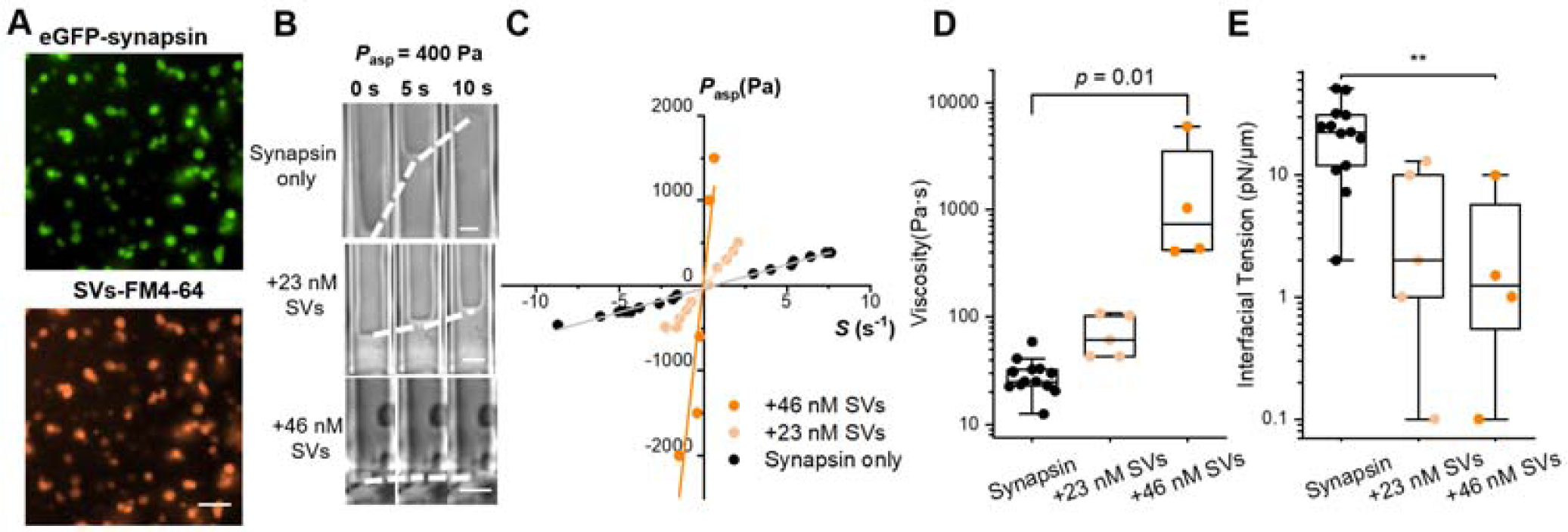
SVs increase the viscosity but lower the interfacial tension of full length synapsin condensates. **(A)** Colocalization of eGFP-synapsin (green) and SVs labeled with FM^TM^ 4-64 (orange). Scale bar, 10 μm. **(B)** Time-lapse transmitted light images showing the deformation of synapsin condensates with increasing concentration of SVs. Dashed lines trace the change in *L*_p_ under 400 Pa aspiration pressure. Scale bars, 2 μm. **(C)** Relation between aspiration pressure and shear rate measured on condensates of synapsin only (black), synapsin + 23 nM SVs (light orange), and synapsin + 46 nM SVs (orange). Effect of SV on the viscosity **(D)** and interfacial tension **(E)** of synapsin condensates. All measurements were done with 9 μM FL synapsin in 3% PEG, 150 mM NaCl, 25 mM Tris-HCl, 0.5 mM TCEP, pH 7.4. *p* values were determined using one-way ANOVA followed by *post hoc* Tukey’s test. \*\**p* < 0.01.

## Discussion

MAPAC allowed us to carry out the first direct quantitative study of the material properties of condensates in living cells. We found the viscoelasticity of cellular synapsin condensates spanned over 4 orders of magnitude (Figure 2) but is highly correlated with the partitioning of α-synuclein (Figure 3). Coordinated with in vitro reconstitution (Figure 5), we identified a critical role of α-synuclein in directly regulating the material properties of synapsin condensates (Figure 7A). Our results opened the door to understanding α-synuclein’s roles in regulating the dynamics of synaptic vesicle release ^42, 68^ and in mediating the formation of solid aggregates that are relevant to neurodegenerative diseases (e.g., Lewy bodies in Parkinson’s disease) ^60, 61, 69, 70^.

**Figure 7:**
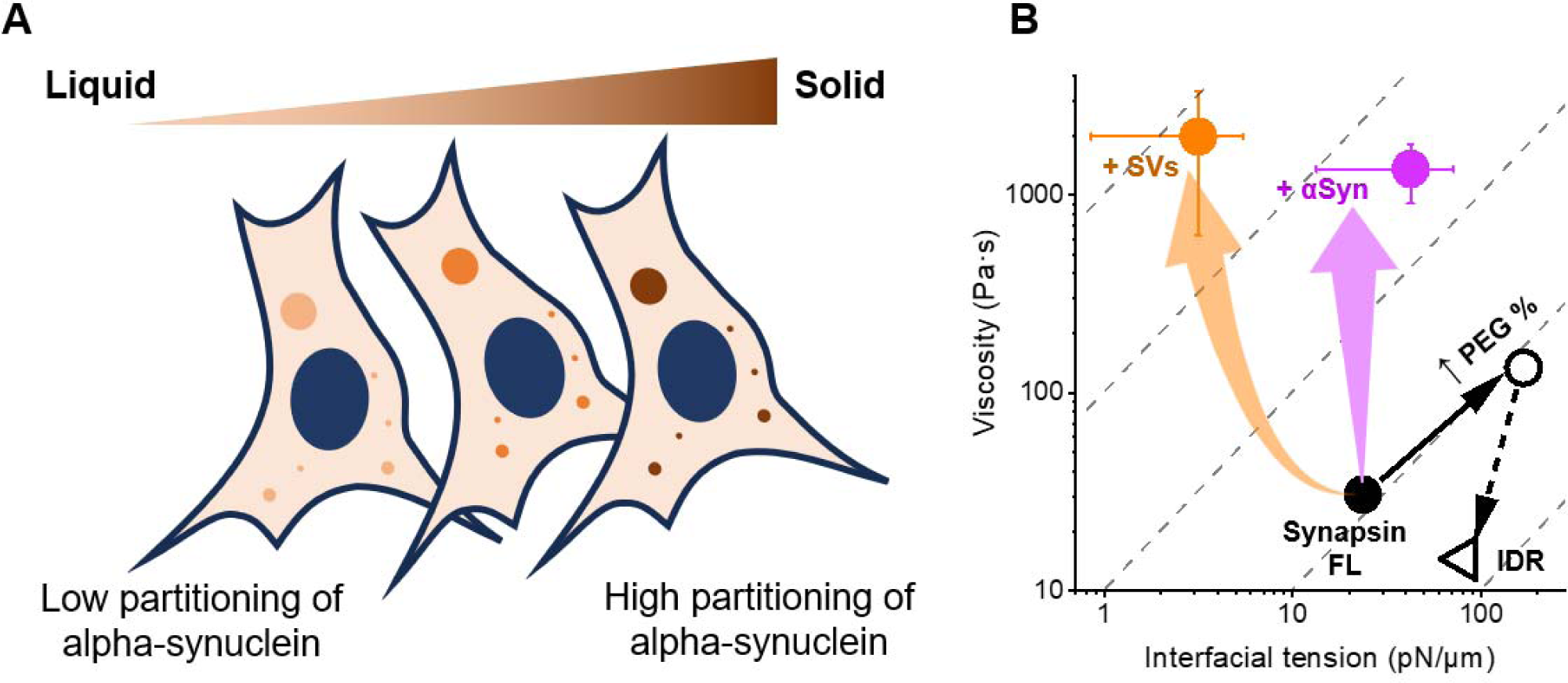
The regulation of synapsin condensate material properties in vitro and in live cells. (A) An illustration showing that the partitioning of α-synuclein dictates the material state of synapsin condensates in cells. (B) Viscosity and interfacial tension of in vitro synapsin condensates. Closed circles represent FL only (black), FL + 9 μM α-synuclein (magenta), or FL + 46 nM SVs (orange) in 3% PEG. Open markers represent FL (circle) and IDR (triangle) in 10% PEG. Error bars are SEM. Dashed lines correspond to constant capillary velocity (i.e., interfacial tension ∼ viscosity).

Cellular synapsin condensates exhibit significantly higher elasticity than those reconstituted in vitro (Figure 5H). The elasticity of typical cellular synapsin/α-synuclein condensates resemble those of hydrogels formed by FUS and hnRNPA (∼ 10^4^ Pa) ^71^, with the lowest elasticity values corresponding to purely liquid droplets and the highest elasticity values approaching those of rubber-like elastomeric proteins such as spider silk (> 10^6^ Pa) ^72^ and amyloid fibers of α-synuclein (∼ 10^9^ Pa) ^73^. Notably, the elasticity of cellular synapsin condensates (Figure 2E) span a similar range as the stiffness of extracellular matrices (soft matrix ∼ 10^2^ Pa, stiff matrix > 10^4^ Pa) that can guide cell migration and differentiation ^74^, suggesting a broad mechanobiological role of biomolecular condensates.

Our measurements of cellular synapsin condensates also show that the morphology of condensates (such as sphericity ^75^) carries little information about their material properties. Although the majority of synapsin condensates were spherical, their viscoelasticity varied drastically, from low-viscosity liquid that allows dynamic molecular exchange to apparent solid that resembles disease-associated inclusions such as Lewy bodies ^60, 61, 69^ (Figure 3). Unlike condensate morphology, the partitioning of key regulatory proteins (i.e., α-synuclein) can be an accurate predictor of condensate material properties (Figure 7A). Posttranslational modifications such as phosphorylation may provide an additional layer of viscoelastic regulation. This is particularly relevant in the case of synaptic activity, where upon activation a global series of phosphorylation events targets numerous proteins ^76, 77^, including synapsin-1 and α-synuclein ^78, 79^. These phosphorylation events may affect the clustering of synaptic vesicles in vertebrate synapses ^40, 79, 80^, further emphasizing the need for methods that allow direct in vivo quantification of viscoelasticity. To fully understand synapsin biology, future efforts need to focus on exploring the roles of synaptic vesicles, disease-related α-synuclein mutations, posttranslational modifications, and other cellular components in modulating the material properties of synapsin condensates under physiological or pathological conditions.

The liquidity of synapsin condensates can be fully controlled via biologically relevant factors in vitro. The presence of α-synuclein, SV, and PEG (mimicking the crowding condition of the cytoplasm) all render synapsin condensates more viscous while removing the N-terminal region of synapsin reduces condensate viscosity (Figure 7B). Viscosity is a kinetic parameter that is governed by the dynamics of molecular interactions within the condensate ^7^, whereas interfacial tension is a thermodynamic parameter that depends on both the dense and the dilute phases ^13^. Indeed, while factors that promote the phase separation of synapsin condensates generally increase condensate viscosity, they can have distinct effects on the interfacial tension of condensates: PEG increases, whereas SVs decrease the interfacial tension of synapsin condensates (Figure 7B), highlighting the complexity of condensate interfaces ^81^. The reduction of interfacial tension by SVs is in line with the effect of protein clusters that serve as Pickering agents ^20^ and with recent observations that lipid vesicles are prone to perturb the interface of condensates ^82^.

We did not observe statistical difference between the interfacial tension of synapsin/α-synuclein condensates in vitro and in cells (Figure 5E, Figure S5). However, the large variation and ∼10-fold higher mean value of the interfacial tension of cellular condensates may reflect biomolecular assemblies (e.g., cytoskeleton) at the condensate interface that resisted the initial condensate deformation under MPA ^83^. Notably, a higher interfacial tension of cellular condensates may also suggest that the intracellular environment is more crowded than the reconstituted condition in vitro (3% PEG for synapsin/ α-synuclein condensates; Figure 4H).

While the viscosity of cellular condensates can be estimated via molecular diffusion measurements (e.g., FRAP, FCS, single molecule tracking), assumptions of molecular size, shape, and boundary conditions are needed to convert the measured diffusion coefficient to a nanoscale viscosity of the condensate ^84^. More importantly, viscosities at the nanoscale often significantly differ from the condensate’s bulk rheology ^12, 85^. The MAPAC platform described here directly measures the overall viscoelasticity of single condensates in their native surroundings. MAPAC shares a large part of its hardware with patch-clamp, a well-established tool in electrophysiology, making it easily accessible to neuroscience, biophysics, and cell biology communities ^49, 86^. Moreover, the electrical recording capacity of MAPAC opens the door to scrutinizing the coupling between electrical, mechanical, and chemical properties of condensates in live cells ^52–54^. Understanding the electrical signals associated with synapsin/α-synuclein condensates (Figure 1) will be a steppingstone towards these directions. Therefore, we expect MAPAC to be a widely accessible and powerful tool for quantitative understanding of a broad range of biomolecular condensates in living organisms, from p granules, stress granules, to those formed by proteins that represent hallmarks of neurodegeneration ^4, 5, 8^.

Our study of synapsin/α-synuclein condensates were carried out in a minimal system in vitro and with ectopically generated condensates in live cells. To better understand the physiological functions of these condensates, future efforts will need to focus on measurements in neuronal synapses in vivo. The typical opening diameter of micropipettes used in MAPAC is 0.5 ∼ 1 μm, best suited for studying cytosolic condensates with a diameter >1.5 μm. While micrometer-sized condensates are widely observed in the literature (Table S1), quantifications of submicron or nuclear condensates in live cells are still challenging. Unlike in vitro quantifications, the interfacial tension of synapsin/α-synuclein condensates in cells showed large measurement uncertainties (Figure 2H, S5). This is partly due to the low resolution of the pressure system used for cellular measurements (resolution ∼100 Pa in cell vs. ∼1 Pa in vitro). The compromise in resolution was needed to reach sufficiently large aspiration pressure to break into the cell and to deform the highly viscoelastic condensates (Methods). Therefore, further developments should focus on nanoscale techniques that can easily penetrate the cell surface and supply large pressure (>10 kPa) with high precision (< 1Pa).

## Supporting information

Supplementary Figures, Supplementary Table, Supplementary Discussions

## Methods

### Cell culture and transfection

HEK 293T cells (ATCC) were grown in Dulbecco’s Modified Eagle Medium (11995065, Thermo Fisher) enriched with 10% Fetal Bovine Serum (FB12999102, Fisherbrand), 1% Penicillin-Streptomycin (15140122, Thermo Fisher). Approximately 1 million cells were seeded into a 100 mm plastic dish. The cells were maintained in an environment at 37°C, with 5 – 10 % CO^2^, and 100% relative humidity.

300,000 cells were seeded onto a 35 mm glass bottom dish (D35-20-1.5-N, Cellvis) precoated with Matrigel Matrix (47743-715-EA, Corning Life Sciences) for transfection. 24 hours after seeding the cell, a mixture of 250µL Opti-MEM, 5 µL P3000 reagent, 7.5 µL Lipofectamine reagent, 1.5 µg of mCherry-Synapsin 1 plasmid ^33^, and 1 µg α-synuclein-Halo ^87^ or α-synuclein-BFP ^33^ plasmid was added to the dish. Imaging and MAPAC experiments were done 24 hours after transfection. Prior to imaging, Halo-α-synuclein expressing cells were incubated with 1 µM JFX-646 Halo Tag Ligand ^88^ for 15 min, and the cell culture medium was replaced with an extracellular imaging buffer (XC buffer). The XC buffer contained 140 mM NaCl, 5 mM KCl, 10 mM HEPES, 10 mM Glucose, 2mM MgCl2, and 2 mM CaCl2, at a pH of 7.4. All chemicals were ordered from Sigma.

### MAPAC experiments

Different from MPA measurements in vitro, micropipettes used in MAPAC were pulled into a cylindrical tip with an opening diameter of 0.5 ∼ 1 μm (PUL-1000, World Precision Instruments). The tip was then bent to an angle of approximately 40° using a microforge (DMF1000, World Precision Instruments). The micropipette was filled with an intracellular buffer (126 mM K-Gluconate, 4 mM KCl, 10 mM HEPES, 2 mM Mg-ATP, 0.3 mM Na2-GTP, 10 mM Phosphocreatine, final pH 7.2, osmolarity 270 – 290 mOsm; all chemicals were ordered form Sigma) and mounted to a headstage that is connected to an Axon 700B amplifier (Molecular Devices) and to a home-made pressure recording device using an Arduino(UNO R3,ELEGOO) controlled pressure sensor (B07N8SX347, FTVOGUE, resolution 100 Pa). The micropipette was then inserted into a dish with HEK 293T cells expressing fluorescently labeled condensates under the voltage clamp mode. Typical resistance of the micropipette was 50 MΩ. A small ejection pressure (∼ −1 kPa) was applied to the micropipette until the pipette tip reached the targeted cell. Then a suction pressure was maintained in the micropipette to form a giga seal near a targeted condensate. Next, the electrical recording was switched to current clamp mode (with I = 0) and sudden suction pressures pulses >10 kPa were applied to break into the cell. After successfully broke into the cell, the resting membrane voltage was recorded. On healthy HEK 293T cells, reported resting membrane voltage are around −40 mV ^51^. To characterize condensate properties under whole-cell patch clamp mode, suction pressures were applied so that the nearby condensate could flow into the tip of the micropipette for subsequent measurements of condensate material properties. Liquid junction potentials were not compensated for in reported cell membrane potential measurements. The voltage recordings were processed through a 2 kHz filter, digitized at a rate of 10 kHz, and collected using Clampex 10.2 software (Molecular Devices).

### Protein expression and purification

#### eGFP-synapsin 1 and eGFP-synapsin 1 IDR

Both proteins were expressed in Expi293F™cells (Thermo Fisher Scientific) for three days following enhancement and was purified as described previously ^26, 33^. In short, cells were lysed in a buffer that contained 25 mM Tris-HCl (pH 7.4), 300 mM NaCl, 0.5 mM TCEP (buffer A) supplemented with EDTA-free Roche Complete protease inhibitors, 25 mM imidazole, 10 µg/mL DNase I and 1 mM MgCl_2_. All purification steps were carried out at 4°C. Debris was removed by centrifugation for 1 hour at 20,000 x g. For affinity purification, soluble supernatant was applied on a Ni-NTA column (HisTrap™HP, Cytiva, ÄKTA pure 25M) for binding. After a wash step (buffer A with 40 mM imidazole) and elution (buffer A with 400 mM imidazole), proteins were concentrated and subjected to size exclusion chromatography (Superdex™ 200 Increase 10/300, GE Healthcare, ÄKTA pure 25M) in 25 mM Tris-HCl (pH 7.4), 150 mM NaCl, 0.5 mM TCEP. Proteins were snap-frozen in liquid nitrogen and stored at-80 °C until further use.

#### α-Synuclein WT and α-synuclein 140C

For condensate partitioning experiments (Figure 5F, 5G, and S9), untagged human α-synuclein and α-synuclein-A140C were expressed from pET28a vector (Novagen). In brief, a 1 L expression culture (LB; 10 g/L tryptone, 5 g/L yeast extract, 5 g/L NaCl) supplemented with 50 µg/mL kanamycin was inoculated with an overnight preculture to achieve a final OD600 of 0.1. Cells were incubated at 37°C and protein expression was induced at an OD600 of 0.7-0.8 with 1 mM isopropylthio-β-galactoside (IPTG). Cells were shifted to 18°C and incubated overnight while shaking. Upon expression, cells were harvested by centrifugation (7,000xg for 15 min at 4°C).

For α-synuclein purification, the pellet was resuspended in 20 mL ice-cold PBS (Gibco 10010015) supplemented with protease inhibitors (Complete EDTA-free, Roche) and a spatula tip of lysozyme (A3711, AppliChem). Suspension was subjected to lysis in a French press. The lysate was cleared by centrifugation (30 min at 50,000xg at 4°C). Nucleic acids were precipitated by adding streptomycin sulfate (S6501, Sigma) to 10 mg/mL final concentration with constant agitation for 30 minutes at room temperature.

For purification of α-synuclein-A140C variant 1 mM DTT was added. Subsequently the nucleic acids were pelleted by centrifugation (30 min at 50,000 x g and at 4 °C). The supernatant was transferred to a clean glass beaker and proteins were precipitated by slowly adding ammonium sulfate to a final concentration of 0.36 g/mL with constant agitation for 60 min at 4 °C. Proteins were collected by centrifugation for 30 min at 50,000 x g (4 °C). The pellet was resuspended in 10 mL PBS and boiled for 20 min. After centrifugation (30 min at 50,000 x g, 4 °C), the α-synuclein containing supernatant was subjected for dialysis against 5 L of 15 mM Tris base solution overnight (132655T, Spectrum™ Labs Spectra/Por™1 6-8 kDa MWCO Standard RC). For dialysis and subsequent ion exchange chromatography of α-synuclein-A140C variant, 1 mM DTT (final concentration) was used in all steps. The dialyzed protein solution was subjected to anion exchange chromatography (HiPrep Q FF 16/10, Cytiva, ÄKTA pure 25 M) in 25 mM Tris-HCl pH 7.7 (Buffer IEX). Elution was performed stepwise against 25 mM Tris-HCl pH 7.7 containing 500 mM NaCl (IEX-HIGH). First, unwanted material was washed out with 30% IEX-HIGH. Second, α-synuclein was eluted with 50% IEX-HIGH (= 250 mM NaCl). Elution fractions were analyzed by SDS-PAGE (18% gel). Fractions containing α-synuclein were concentrated (Amicon® Ultra 4 MWCo 3kD, Merck Millipore) and subjected to size exclusion chromatography (Superdex™ 200 Increase 10/300, GE Healthcare, ÄKTA pure 25M) in 25 mM Tris-HCl (pH 7.4), 150 mM NaCl, 0.5 mM TCEP. α-synuclein concentration was determined at 280 nm by using a molar extinction coefficient of 5960 M^−1^ cm^−1^. Proteins were snap-frozen in liquid nitrogen and stored at −80 °C until further use.

For MPA measurements (Figure 5A – 5E, S8), α-synuclein was expressed using the pT7-7 aSyn WT vector (Addgene #36046) in BL21(DE3) bacteria (NEB). Briefly, 50mL starter cultures of LB media (Fisher Scientific, BP1426-2) with 50mg/mL ampicillin (Fisher Scientific, BP1760), were inoculated with freshly picked colonies and grown overnight (37°C, 160RPM). The starter cultures were used to inoculate 1L flasks of fresh LB with ampicillin and grown at 37°C (180RPM) until the OD600 measured 0.6-0.8. Expression was induced with 1mM isopropylthio-β-galactoside (IPTG; Teknova I3325) and grown in the incubator overnight (20°C, 160RPM), after which cells were subsequently harvested by centrifugation (4000 rpm, 4°C, 30 minutes) and stored at −80°C until use.

α-Synuclein-A140C was generated from the pT7-7 aSyn WT vector using the Q5 Site-Directed Mutagenesis Kit (NEB, E0554) according to manufacturer’s instructions. Primers were designed using the NEBaseChanger tool and the sequences used were as follows: 5’-CGAACCTGAATGCTAAGAAATATCTTTGC-3’ and 5’-TAGTCTTGATACCCTTCC-3’.

α-Synuclein purification was accomplished by suspension of the bacterial pellet in fresh PBS (Fisher Scientific, BP2944), followed by sonication on ice (15 sec on, 30 sec off; 8 cycles total). The solution was then boiled, followed by centrifugation to clear the lysate (20,000 rpm, 4 °C, 30 minutes). Streptomycin sulfate (Fisher Scientific, BP910-50) was added to the supernatant to a final concentration of 10mg/mL and mixed at 4°C for 30 minutes, followed by centrifugation using the same conditions as before. The protein was precipitated from the supernatant by adding ammonium sulfate (Fisher Scientific, BP212-212) to a final concentration of 0.361G/mL and mixing at 4°C for 1 hour. The protein was pelleted by centrifugation, and the protein pellet was resuspended in buffer A (32.2mM Tris Base, pH 7.8). α-Synuclein was purified via FPLC (Cytiva A□KTA Pure) using HiTrap Q HP anion exchange columns (Cytiva) and eluted at 50% buffer B (32.2mM Tris Base, 500mM NaCl, pH 7.8). Final proteins were aliquoted, flash frozen in liquid nitrogen, and stored at −80°C until use. α-Synuclein A140C was purified in the same way as described above, but with the addition of 1mM DTT.

To label α-synuclein-A140C, first, the protein solution was centrifuged at 16,000 g for 30 minutes with a 3 kD cut-off filter (Amicon Ultra Centrifugal Filter, 3 kDa MWCO, Merck Millipore) to remove previous buffer. The centrifugation step was repeated, when necessary, to reach a targeted protein concentration of 2 mg/ml in PBS. Next, the protein solution was centrifuged at 16, 000 g for 10 minutes with a 100 kD cut-off filter (Amicon Ultra Centrifugal Filter, 100 kDa MWCO, Merck Millipore) to remove potential protein aggregations. Then, the protein solution was incubated with 2x excess Alexa Fluor™ 647 C2 Maleimide (Invitrogen™) for 2 h at room temperature. Next, the protein dye mixture was passed through a desalting column (Zeba Spin Desalting Columns, Thermo Scientific) to remove excess dye. The concentration of labelled protein and degree of labelling were determined by Nanodrop (Invitrogen™ Nanodrop™ One, Fisher scientific).

### Preparation of synaptic vesicles

Synaptic vesicles (SVs) were isolated according to previous publications ^33^. Briefly, 20 rat brains were homogenized in ice-cold sucrose buffer (320 mM sucrose, 4 mM HEPES-KOH, pH 7.4 supplemented with 0.2 mM phenylmethylsulfonylfluoride and 1 mg/ml pepstatin A). Debris was removed by centrifugation (10 min at 900 gAv, 4 °C) and the resulting supernatant was further centrifuged (10 min at 12,000 gAv, 4 °C). The pellet containing synaptosomes was washed once, and synaptosomes were lysed by hypo-osmotic shock. Free, released SVs were obtained after centrifugation of the lysate for 20 min at 20,000 gAv, 4 °C. The supernatant containing the SVs was further ultracentrifuged for 2 h at 230,000 gAv, resulting in a crude synaptic vesicle pellet. Subsequently, SVs were purified by resuspending the pellet in 40 mM sucrose followed by centrifugation for 3 h at 110,880 gAv on a continuous sucrose density gradient (50–800 mM sucrose). SVs were collected from the gradient and loaded to size-exclusion chromatography on glass beads (300 nm diameter), equilibrated in glycine buffer (300 mM glycine, 5 mM HEPES, pH 7.40, adjusted using KOH). This allows for the separation of synaptic vesicles from residual larger membrane contaminants. SVs were pelleted by centrifugation for 2 h at 230,000 gAv and resuspended in sucrose buffer by homogenization before being aliquoted into single-use fractions and snap-frozen in liquid nitrogen until use.

All animal experiments were approved by the Institutional Animal Welfare Committees of the Charité University Clinic (Berlin, DE) and Max Planck Institute for Biophysical Chemistry (Göttingen, DE).

### Condensate fusion experiments

An optical tweezers-assisted fusion assay was carried out and analyzed following the protocol we previously described ^43^. Two condensates were independently trapped by a pair of separate optical tweezers at minimal power (Tweez305, Aresis, Slovenia; mounted on a Nikon Ti2-A inverted microscope). The fusion events were captured at 20 Hz with a 60x water objective. MATLAB R2020b was used to analyze the captured images. The images were fitted to a Gaussian ellipse. The aspect ratio (*AR*), which is the ratio between the major and minor axes of the ellipse, was plotted over time (Figure 4B). The time of fusion τ_fusion_ was deduced by fitting the relaxation in *AR* to a stretched exponential decay ^89^.

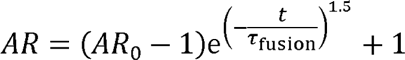

The condensates’ length was designated as the geometric mean of the condensate diameters prior to fusion. The ratio of viscosity over interfacial tension was calculated from the slope of the fusion time vs. length (Figure 4C).

### MPA experiments in vitro

The micropipette aspiration experiments were carried out on a Ti2-A inverted fluorescence microscope (Nikon, Japan) equipped with a motorized stage and two motorized 4-axes micromanipulators (PatchPro-5000, Scientifica) and a multi-trap optical tweezers (Tweez305, Aresis, Slovenia) according to the protocol we reported previously ^43^ with minor modifications.

Micropipettes were made by pulling glass capillaries using a pipette puller (PUL-1000, World Precision Instruments). The pipette tip was then cut to achieve an opening diameter ranging from 2 to 5 μm. Subsequently, the pipette was bent to an angle of approximately 40° using a microforge (DMF1000, World Precision Instruments).

The micropipette was filled with the same buffer as the buffer for synapsin (25 mM Tris-HCl, 150 mM NaCl, 0.5 mM TCEP, pH = 7.4) using a MICROFIL needle (World Precision Instruments). The filled micropipette was then mounted onto a micromanipulator. The rear end of the pipette was connected to an automatic pressure controller (Flow-EZ, Fluigent; Pressure resolution 1 Pa). The MPA experiments were conducted in glass-bottom dishes (D35-20-1.5-N, Cellvis), under brightfield illumination to minimize potential artifacts associated with fluorescence excitation. Fluorescence images were taken after MPA experiment to confirm the protein species. Typically, optical tweezers-assisted condensate fusion experiments were first carried out to achieve a large (> 5 μm) condensate for easier MPA measurements and analysis. A secondary micropipette was used to hold the condensate during MPA. To minimize sample evaporation, 1.5 mL Milli-Q water was added to the edge of the dish, and the dishes were covered with a thin plastic wrap with a ∼2 mm hole for pipette insertion.

We observed that in vitro synapsin condensates always wet the inner wall of uncoated micropipettes. Therefore, the analysis of the MPA data follows the protocol described in reference ^44^. Briefly, normalized aspiration length (*L*_p_/*R*_p_) was segmented according to the pressure steps. For each segment, the slope of a linear fitting of (*L*_p_/*R*_p_)^2^ vs. time gives the effective shear rate *S*. Then the slope of *P*_asp_ vs. *S* gives the 4η, the intercept gives 2γ/*R*_p_.

### Condensate partitioning measurement

For in vitro partitioning measurements (Figure 5F,5G, and S9), eGFP-synapsin 1, Polyethylene glycol 8,000 (PEG 8,000), sample buffer, and (if required) maleimide-labeled α-synuclein(A140C)-647 were mixed in a reaction tube (10 µl final volume). From there, the sample was pipetted onto a glass bottom microscopy dish for imaging immediately. The sample solution consisted of 5 µM eGFP-synapsin 1, 3% PEG 8,000, 150 mM NaCl, 25 mM Tris-HCl, 0.5 mM TCEP, pH 7.4 with varying concentrations of α-synuclein(A140C)-647 as indicated in the text and the corresponding figures. Each condition was assessed in at least three independent reconstitutions.

The microscopy data was acquired at the Advanced Biomedical Imaging (AMBIO) Facility at Charité Medical Center, using the Eclipse Ti Nikon Spinning Disk Confocal CSU-X, equipped with 2 EM-CCD cameras (AndorR iXon 888-U3 ultra EM-CCD), Andor Revolution SD System (CSU-X), objectives PL APO 60/1.4NA oil immersion lens. Z-stacks (step-size: 0.3 µm) were acquired with piezo stage z-motor at two different wavelengths (488 nm for eGFP-synapsin 1 and 638 nm for α-synuclein(A140C)-647) after loading the sample onto the surface of the microscopy dish over a duration of 15 minutes over a duration of 15 minutes. Each z-series was imaged twice at two different laser intensities. The lower intensity was always used for the analysis. The Exposure times were set to 200 ms (488 nm) and 100 ms (638 nm), and the EM Gain (30 MHz at 16-bit) Multiplier was set to 300 during all these experiments. NIS Elements 5.21.02 software was used to acquire the microscopy data. Fiji ImageJ (NIH) was used to analyze the data.

For the diameter analysis, the equatorial z-plane of most condensates was selected. Then, the area (µm^2^) of all condensates within 60 µm of the edge of the sample solution was quantified by applying a Gaussian Blur (Sigma: 2.00 px) and generating a binary image using the Otsu algorithm, based on the images from the synapsin channel. Only particles with a circularity ≥ 0.9 were included in the analysis. The diameter of the analyzed condensates was calculated based on the measured area.

For the analysis of the raw and relative fluorescence intensities (partitioning coefficient), a squared section of approximately 40 µm × 40 µm was generated, and the equatorial z-plane was chosen for analysis. Here, the regions of interest (ROIs) were selected by applying a Gaussian blur (Sigma: 2.00 px) and generating a binary image using the Otsu algorithm, based on the images from the synapsin channel, only including particles with a circularity ≥ 0.9 in the analysis. Then, ten additional ROIs of similar size to the condensates were placed in the dilute phase on the same z-plane, and the mean fluorescent intensities for each ROI (both condensates and dilute phase) were measured at both wavelengths. A correction value, accounting for the background fluorescence detected by the camera in the absence of any fluorophore, was subtracted from every readout value, and the mean corrected fluorescence intensities were calculated for both channels in the condensates and the dilute phase. The partitioning coefficients for both proteins were computed by dividing each condensate’s corrected mean fluorescence intensity by the mean corrected fluorescence intensity in the dilute phase for the same corresponding image.

For the partitioning coefficient of cellular condensates, first confocal and widefield fluorescence images of the same condensate-containing HEK 293T cells were taken on a Zeiss Axio Observer 7 inverted microscope equipped with an LSM900 laser scanning confocal module and using a 63×/1.4 NA plan-apochromatic, oil-immersion objective.

The images were then analyzed using ImageJ. As shown in Figure S6a, the condensate partitioning coefficient (ll) is defined as the background-corrected mean fluorescence of in the condensate (*I*_condensate_) divided by the background-corrected mean fluorescence of the surrounding cytosol (*I*_cytosol_) for the studied proteins. The background was measured as the mean extracellular fluorescence.

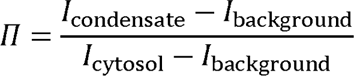

The partitioning coefficients measured with confocal and widefield microscopy for the same condensates were used to establish a conversion function between π_confocal_ and π_widefield_ (Figure S6B). The conversion function (π_confocal_ = 6.7π_widefield_ - 5.7) was used to assign pseudo-confocal partitioning coefficients on condensates measured in MAPAC with a widefield configuration.

#### Acknowledgments

We thank Drs. James Roggeveen, Howard Stone, Eric Dufresne, Conor McClenaghan, Che-Yu Lin, Benjamin Schuster, Cliff Brangwynne, Wilma Olson, Sagar Khare, Mengying Deng, and members of the Rutgers LLPS journal club for helpful discussions. We thank Dr. Benjamin Schuster lab for access to confocal microscopy. We thank Dr. Luke Lavis lab for sharing JFX dyes. The project is supported by the National Institute of General Medical Sciences of the National Institutes of Health (NIH) under Award Number R35GM147027 (Z.S.) and by the National Institute on Drug Abuse, the National Institute of Neurological Disorders and Stroke, and the National Institute of Mental Health of the NIH under Award Number R21DA056322 (Z.S.). D.M. is supported by the start-up funds from DZNE, the research grant from the German Research Foundation (MI 2104), and the European Research Council (MEMLESSINTERFACE, 101078172); views and opinions expressed are however those of the authors only and do not necessarily reflect those of the European Union or the European Research Council Executive Agency; neither the European Union nor the granting authority can be held responsible for them. J. B. acknowledges NIH grant: R35GM136431. Z.P.P. acknowledges NIH grant: R21MH126420. C.H. is supported by a fellowship of the Innovative Minds Program of the German Dementia Association. X.S. is supported by a Predoctoral Fellowship from the Autism Science Foundation. The Child Health Institute of New Jersey is partially funded by the Robert Wood Johnson Foundation (RWJF #74260).

## Author contribution

Z.S., D.M. and Huan W. conceived the idea and designed the experiments. Huan W. performed most of the experiments. Huan W. and Z.S. analyzed the data. C.H. and Han W. purified synapsin proteins, synaptic vesicles, α-synuclein for in vitro partitioning measurements, and cloned plasmids for live cell experiments. J.V.T. and C.H. performed the experiments in Figure 5F, 5G, and S9. Z.P.P. and X.S. guided the electrophysiology experiments. J.B. and J.E. designed and purified α-synuclein proteins for MPA. Huan W., Z.S., C. H., and D.M. wrote the manuscript with input from all authors.

## Competing interests

The authors declare no competing interests.

## Notes

### Competing Interest Statement

The authors have declared no competing interest.

### Summary of Updates

Significantly changed data presentation and discussion.

