## Supplementary Figures, Supplementary Table, Supplementary Discussions for "Live-Cell Quantification Reveals Viscoelastic Regulation of Synapsin Condensates by α-Synuclein"

**Supplement**

Supplementary Figures S1 to S10

Supplementary Movie S1 to S2

Supplementary Table S1

Supplementary Discussions

Supplementary References

**Figure S1**

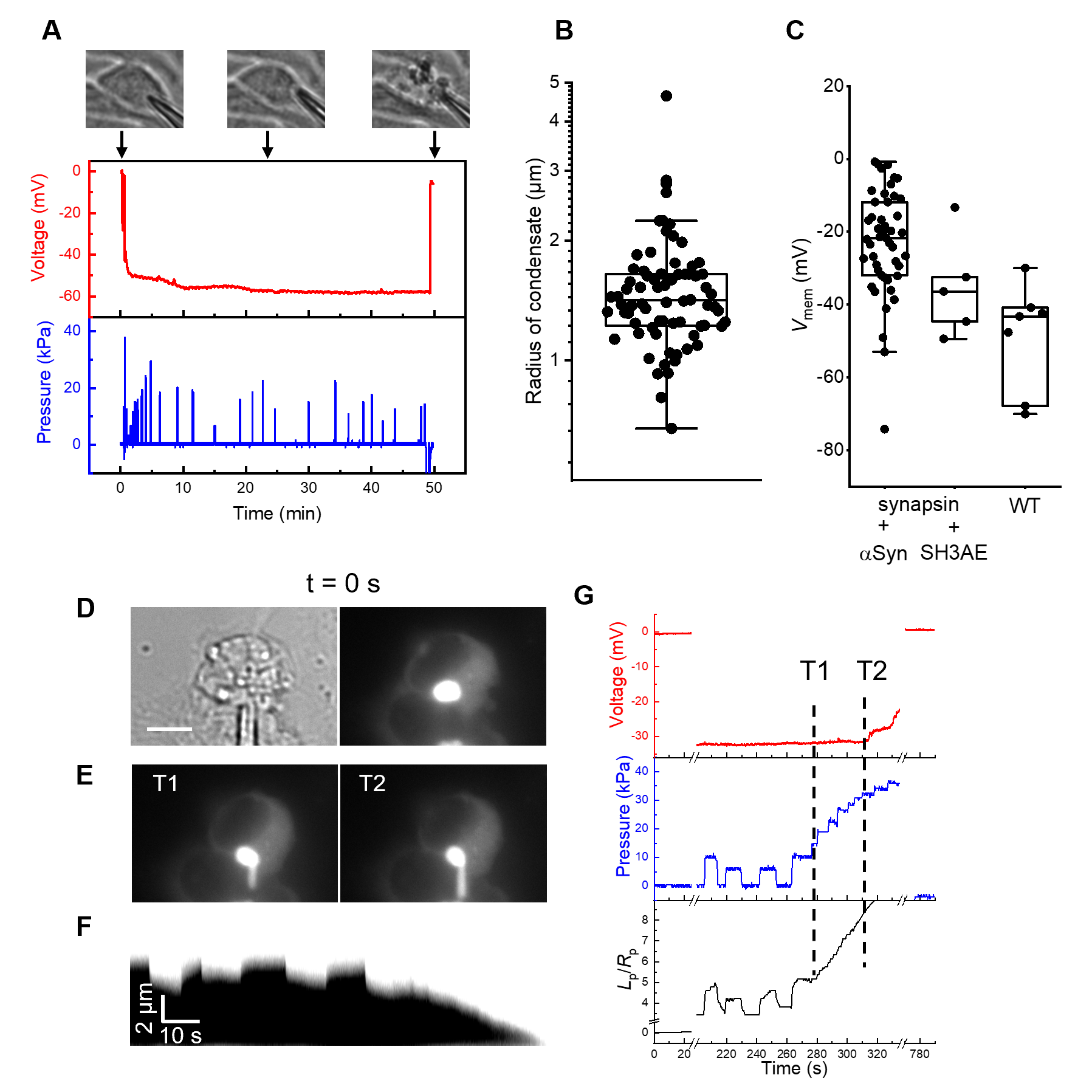

**Figure S1. (A)** Testing of the stability of whole-cell patch clamp in non-transfected HEK 293T cells. **(B)** Condensate radii in HEK 293T cells co-expressing synapsin and α-synuclein. **(C)** Resting cell membrane potential for non-transfected cells, cells transfected with synapsin and α-synuclein, and cells transfected with synapsin and intersectin SH3 domains. **(D)** Transmitted light (left) and fluorescence (right) images of a HEK-293T cell expressing synapsin1-GFP and intersectin-SH3. The cell is under whole-cell patch clamp, but with aspiration pressure *P*_asp_ = 0. **(E)**, Fluorescence images of the cell under *P*_asp_ > 0 (suction), before (T1) and right after (T2) the membrane seal became leaky. We estimate an apparent condensate elasticity ~ 2000 Pa before T1. **(F)** Kymograph of the aspirated part of the condensate from t = 200 to t = 320 s. **(G)** Recordings of membrane voltage (red), *P*_asp_ (blue), and normalized aspiration length (black) during the experiment in **D - F**.

**Figure S2**

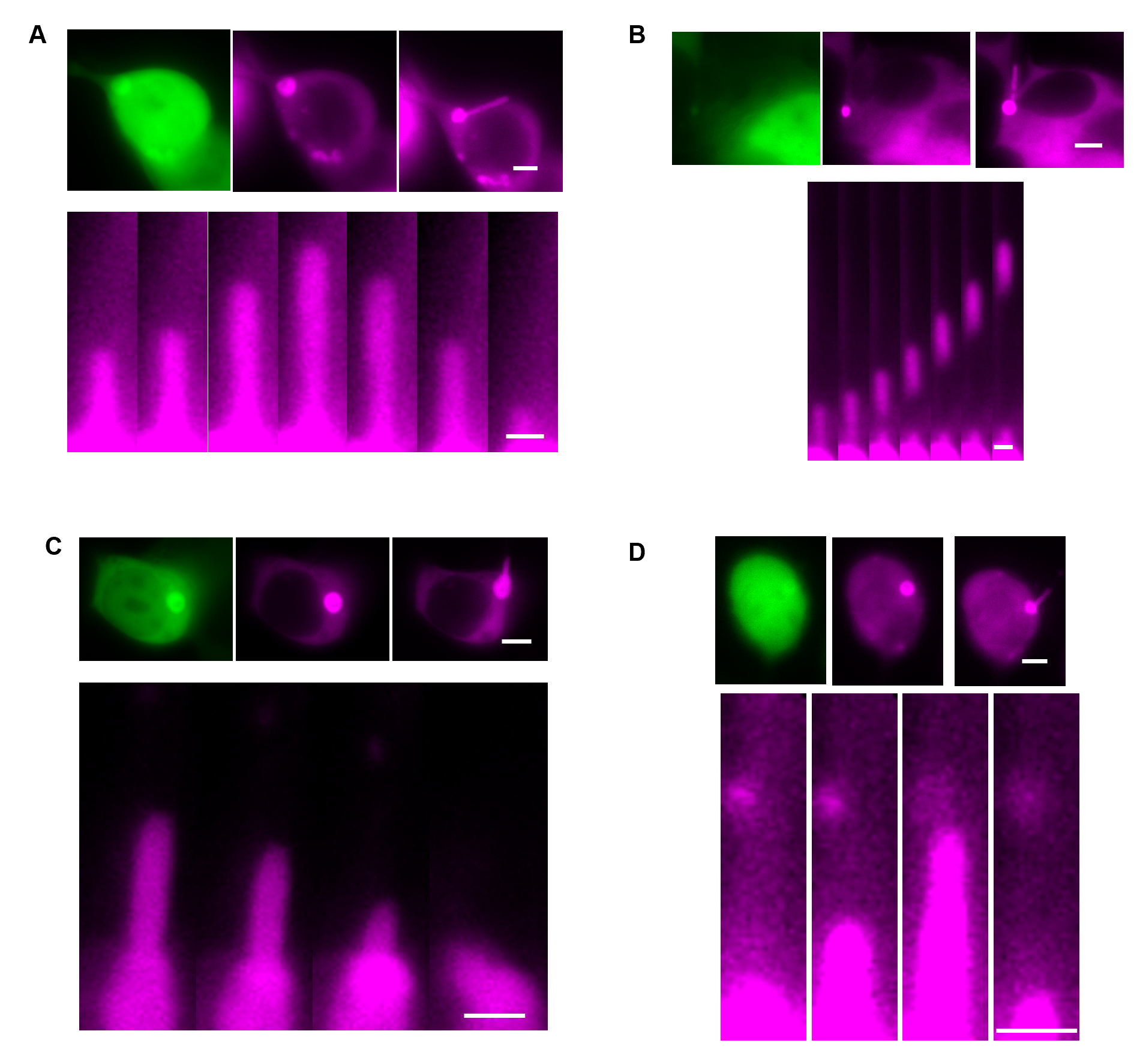

**Figure S2. Cellular synapsin/α-synuclein condensates do not wet the inner wall of micropipettes.**

**(A)** Condensates inside the micropipette formed a convex meniscus. **(B)** Condensates inside the pipette underwent a necking instability and broke apart. **(C, D)** Aspiration of condensates did not leave behind any measurable fluorescence inside the micropipette. In (**A)**- (**D)**, the upper panels show fluorescence images of the entire cell (green: α-synuclein; magenta: synapsin), the condensate was being aspirated in the synapsin channel as shown in the right image. Scale bars, 5 μm. The lower panels show time lapse responses of the aspirated portion of each condensate. Scale bars, 2 μm.

**Figure S3**

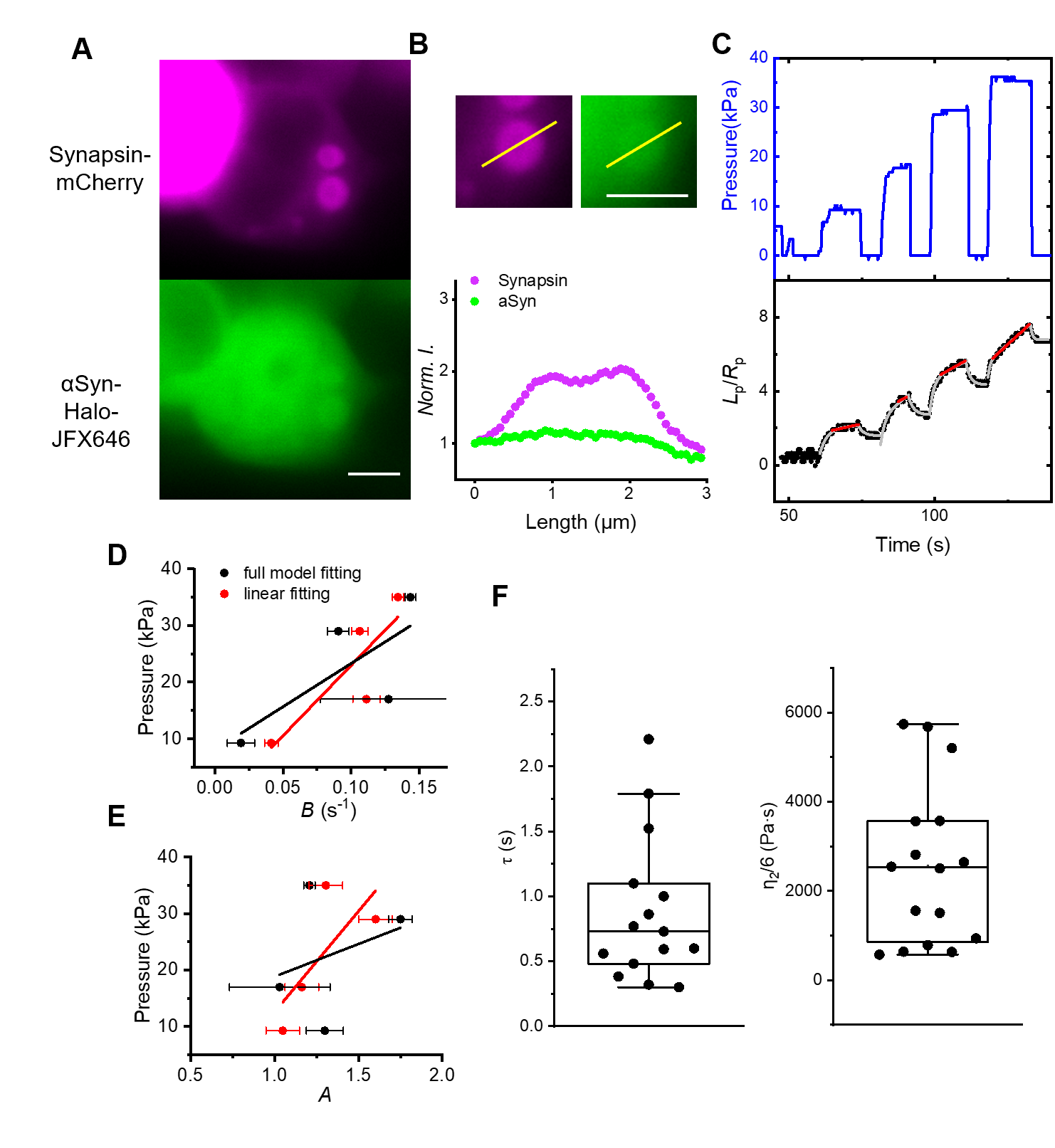

**Figure S3. (A)** Fluorescence images of a HEK 293T cell co-expressing mCherry-synapsin (magenta) and α-synuclein-Halo labelled with JFX-646 (green). **(B)** Intensity profiles across one of the condensates. Scale bar, 5 μm. **(C)** Normalized aspiration length (black), in response to stepwise aspiration pressure (blue). Grey curves are fits to the full model (eq. 1); red lines are linear fits. **(D-E)** Relations between aspiration pressure and long-term shear rate (**D)** and normalized amplitude of elastic response (**E)**. Black and red represent results from the full model and linear model, respectively. Lines are linear fits. **(F)** Value of *η*_2_ (τ × E) determined from fitting to the full model.

**Figure S4**

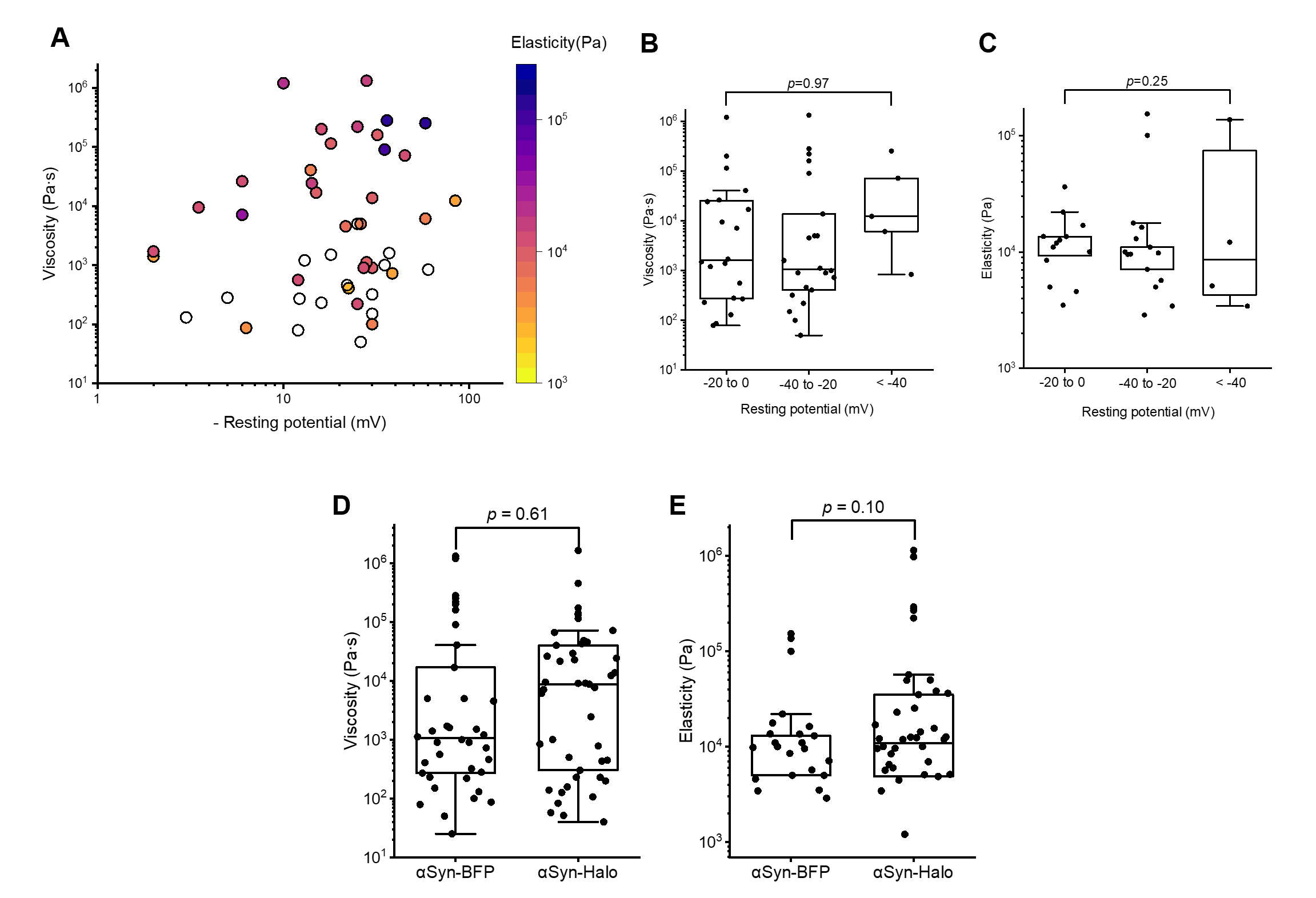

**Figure S4. Measured viscoelasticity of cellular synapsin/α-synuclein condensates is independent the resting potential of the cell membrane and independent of the choice of fluorescent tags.**

**(A)** Condensate viscosity vs. the resting voltage of the cell membrane. Color encodes the elasticity of condensates. Viscosity (**B**) and elasticity (**C**) of condensates grouped according to the resting potential of the cell. *p* values were determined using one-way ANOVA. Measured viscosity (**D**) and elasticity (**E**) of cellular synapsin/α-synuclein condensates is independent the fluorescent tag on α-synuclein. *p* values are from Student’s t-test.

**Figure S5**

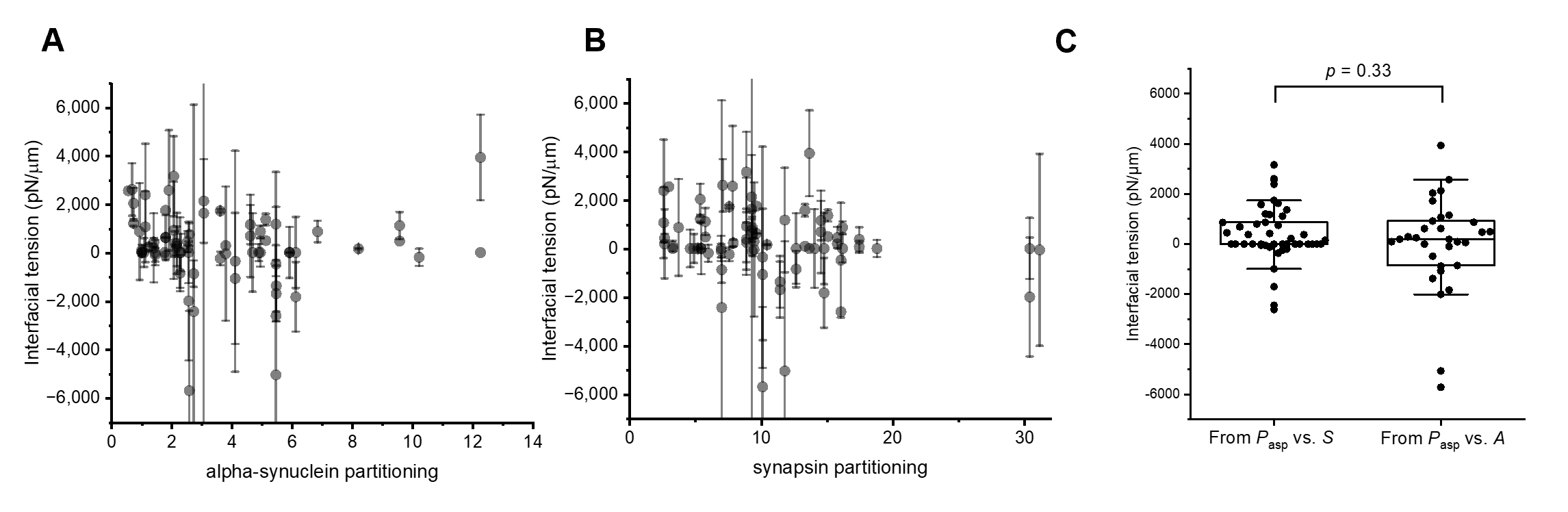

**Figure S5.** Interfacial tension of cellular synapsin/α-synuclein condensates is independent of the partitioning of α-synuclein (**A**), synapsin (**B**), or the model used for data analysis (**C**). Error bars represent the standard deviation in fitting. *p* values are from Student’s t-test.

**Figure S6**

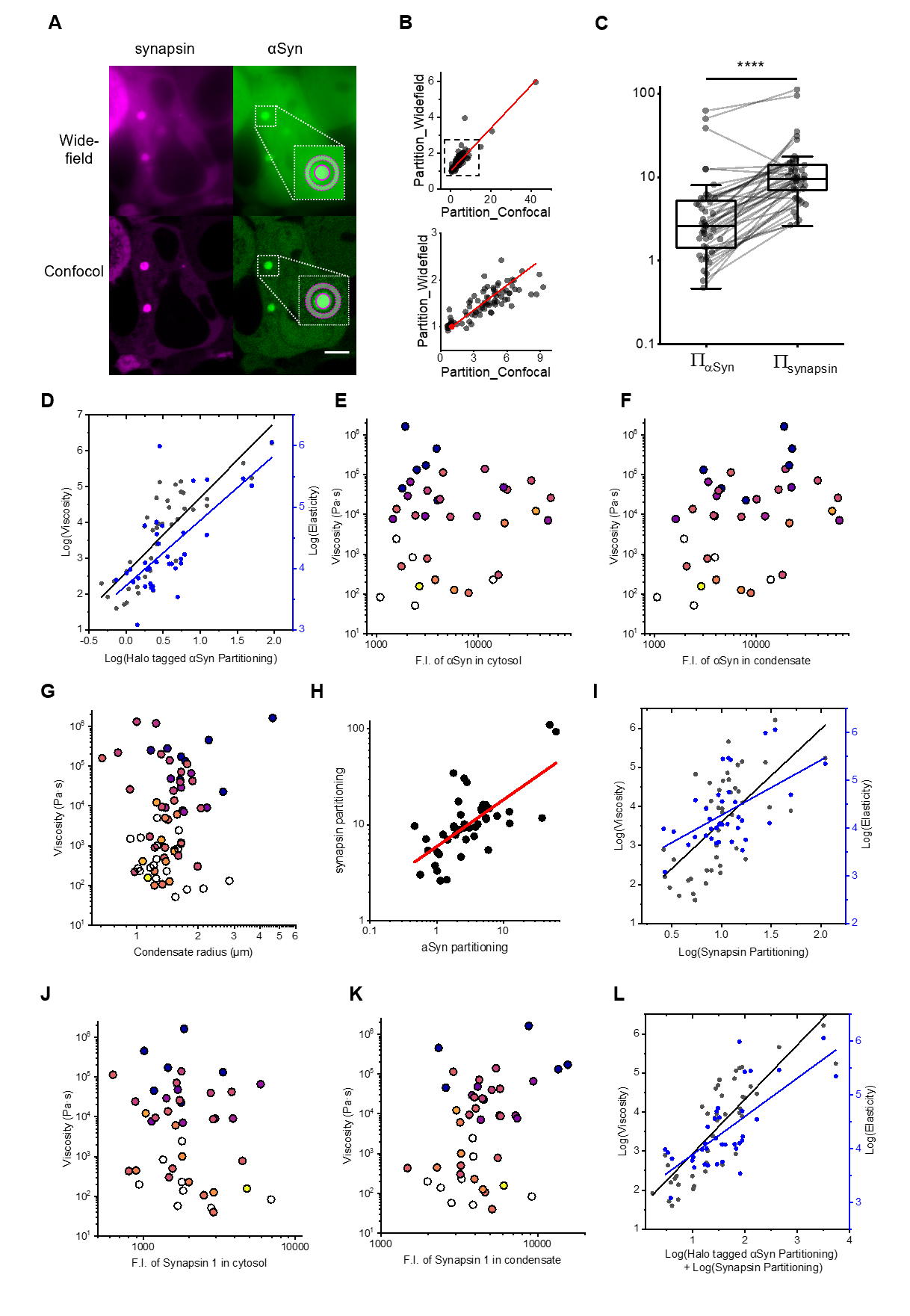

**Figure S6. (A)** Simultaneous widefield (upper) and confocal (lower) fluorescence imaging of HEK 293T cells co-expressing synapsin (magenta) and α-synuclein (green). The solid circle and dashed donut shape represent the ROIs for the condensate and for the dilute phase, respectively. Scale bar, 5 μm. **(B)** Correlation between the apparent partitioning coefficients measured via widefield and the true partitioning coefficients measured via confocal. Line represents a linear fit: $\Pi_{\mathrm{confocal}}=6.7\Pi_{\mathrm{widefield}}-5.7$ (Pearson’s r = 0.93). The red dot represents (1,1). The boxed region in the upper panel is zoomed-in below. **(C)** Comparison between $\Pi_{\alpha Syn}$ and $\Pi_{\mathrm{synapsin}}$ for condensates in Figure 3. **** *p* < 10^-4^, paired t-test. **(D)** Dependence of condensate viscoelasticity on $\Pi_{\alpha Syn}$ (r = 0.84 for viscosity, r = 0.74 for elasticity). **(E)** Dependence of condensate viscoelasticity on the mean fluorescence of the cytosol that represents the concentration of α-synuclein in the dilute phase (r = 0.08). **(F)** Dependence of condensate viscoelasticity on the mean fluorescence of the condensate that represents the concentration of α-synuclein in the dense phase (r = 0.39). **(G)** Dependence of condensate viscoelasticity on condensate radius (r = 0.14). **(H)** Correlation between $\Pi_{\alpha Syn}$ and $\Pi_{\mathrm{synapsin}}$ (r = 0.68). **(I)** Dependence of condensate viscoelasticity on $\Pi_{\mathrm{synapsin}}$ (r = 0.66 for viscosity, r = 0.60 for elasticity). **(J)** Dependence of condensate viscoelasticity on the mean fluorescence of the cytosol that represents the concentration of synapsin in the dilute phase (r = -0.19). **(K)** Dependence of condensate viscoelasticity on the mean fluorescence of the condensate that represents the concentration of synapsin in the dense phase (r = 0.34). **(L)** Dependence of condensate viscoelasticity on $(\Pi_{\alpha Syn}+\Pi_{\mathrm{synapsin}})$ (r = 0.84 for viscosity, r = 0.74 for elasticity).

**Figure S7**

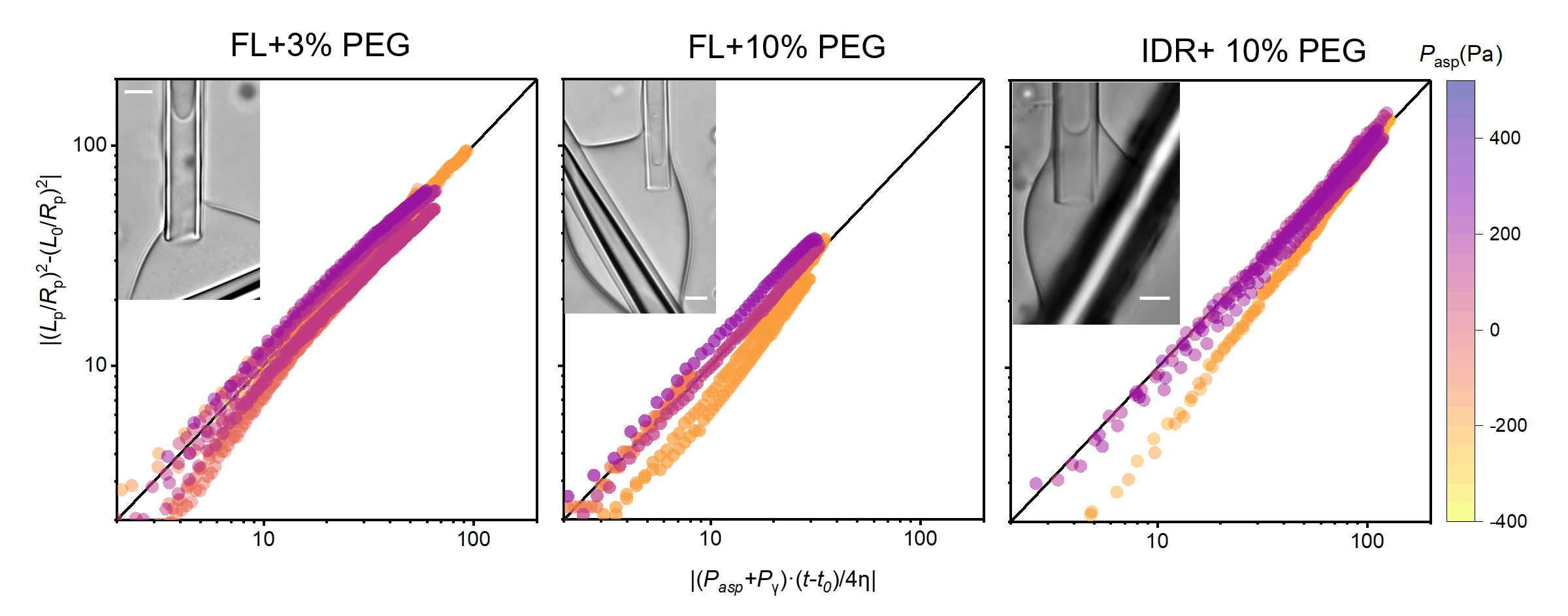

**Figure S7. MPA of synapsin condensates.**

Plots of rescaled and segmented data according to ref^1^. The color of the traces represents applied aspiration pressure. The x axis is rescaled time, and the y axis is shifted (*L*_p_/*R*_p_)^2^. The black lines are y = x. Transmitted light images of micropipette-aspirated synapsin condensates are shown for each condition. Scale bars, 5μm.

**Figure S8**

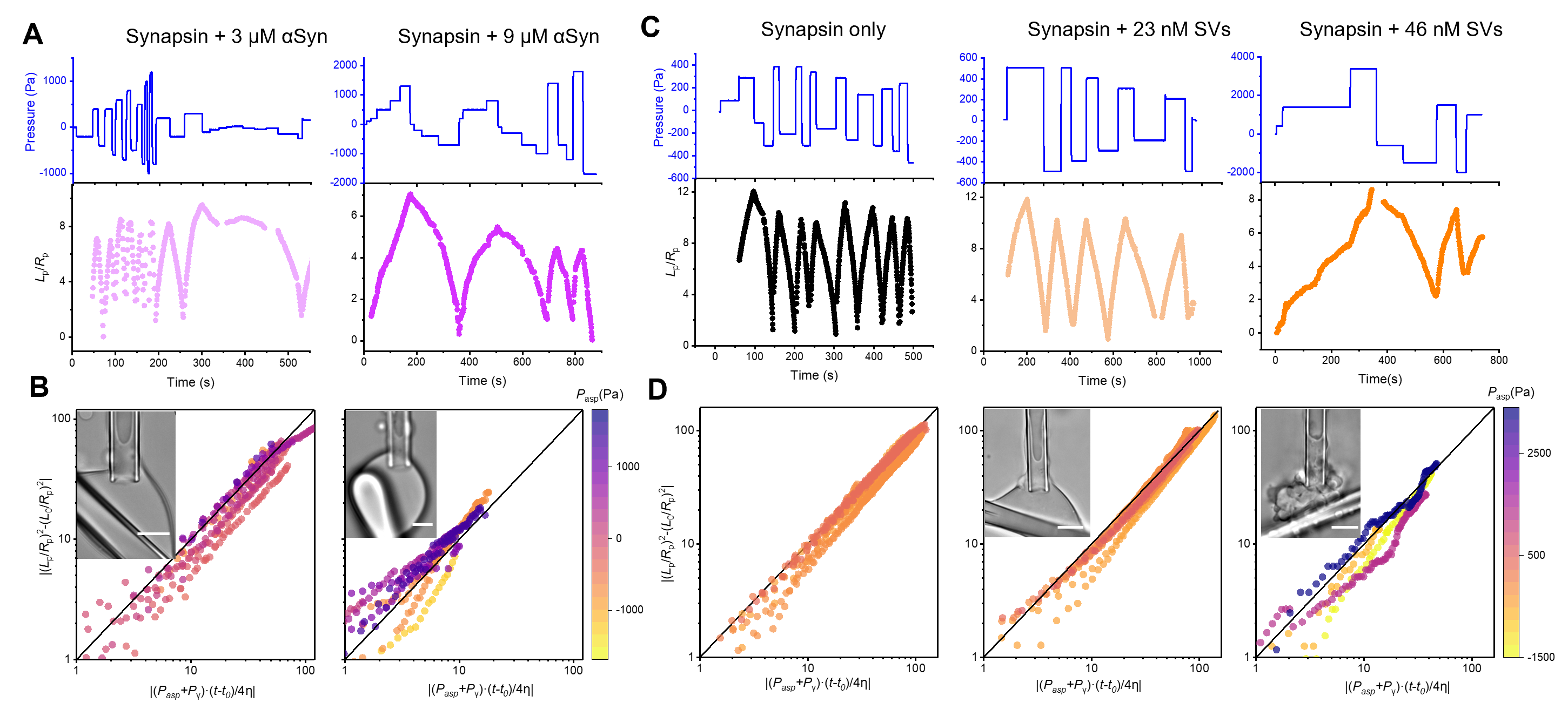

**Figure S8. MPA of synapsin/SV and synapsin/α-synuclein condensates.**

**(A)** Aspiration pressure (upper) and normalized aspiration length (lower) during MPA of synapsin + 3 μM α-synuclein (left), and synapsin + 9 μM α-synuclein (right). **(B)** Same as (B) but plotted for data in (**A)**. **(C)** Aspiration pressure (upper) and normalized aspiration length (lower) during MPA of synapsin only (left), synapsin + 23 nM SVs (middle), and synapsin + 46 nM SVs (right). **(D)** Plots of rescaled and segmented data of (**C)** according to ref^1^. The color of the traces represents applied aspiration pressure. The x axis is rescaled time, and the y axis is shifted (*L*_p_/*R*_p_)^2^. The black lines are y = x. Transmitted light images of micropipette-aspirated synapsin/SV condensates are shown for each condition. All scale bars, 5μm.

**Figure S9**

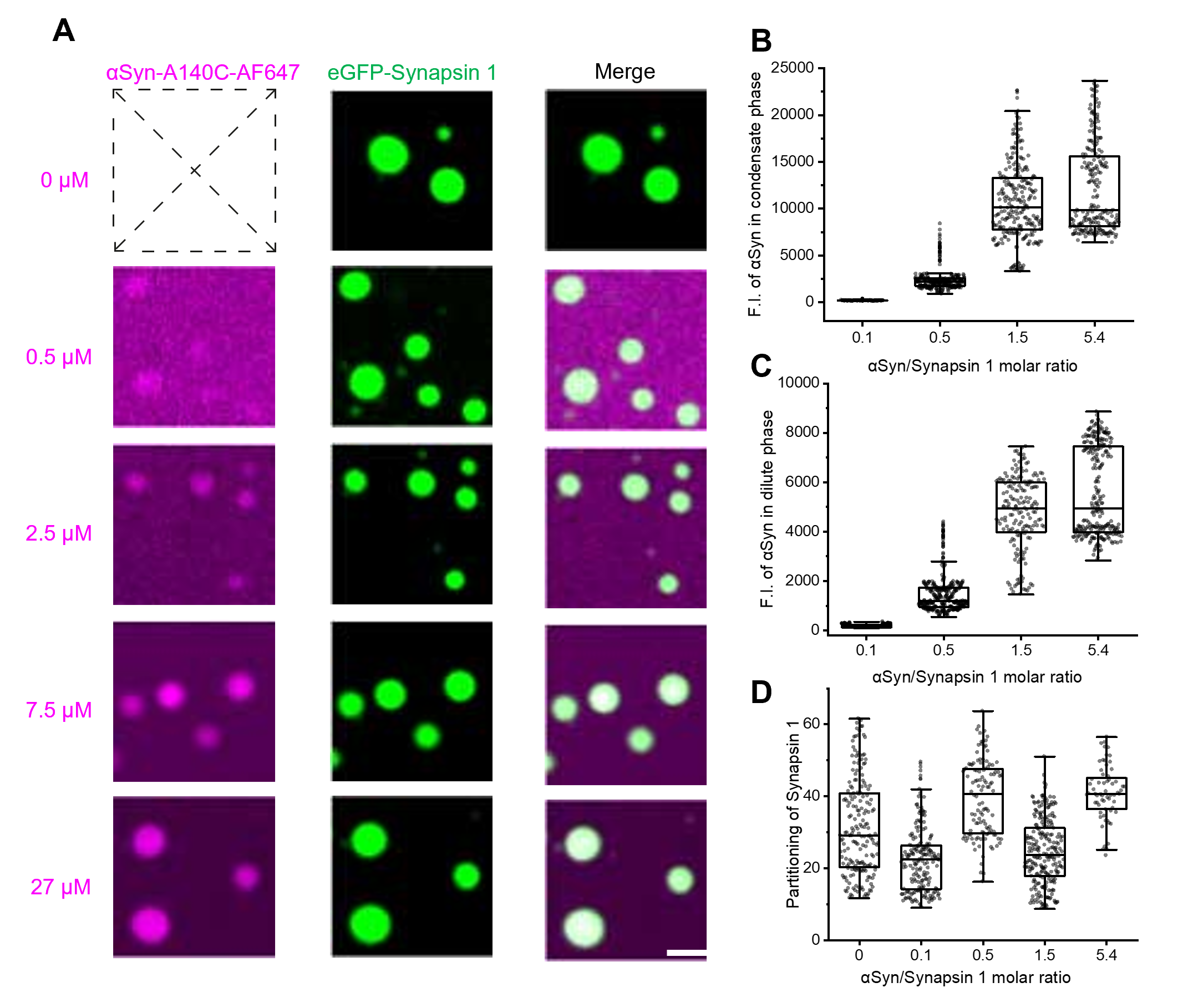

**Figure S9. Condensate partitioning of α-synuclein and synapsin.**

**(A)** Representative confocal microscopy images of in-vitro synapsin/α-synuclein condensates at different concentrations of α-synuclein. All measurements are performed with 5 µM eGFP-synapsin 1 and an increasing amount of α-synuclein (A140C)-Alexa Fluro 647 (top to bottom: no α-synuclein, 0.5 µM, 2.5 µM, 7.5 µM, and 27 µM), all in a buffer containing 150 mM NaCl, 25 mM Tris-HCl, 0.5 mM TCEP, Ph 7.4. Scale bar: 2 μm. **(B)** Effect of α-synuclein concentration on the fluorescent intensity of α-synuclein in synapsin condensate. **(C)** Effect of α-synuclein concentration on the fluorescent intensity of α-synuclein in the dilute phase. **(D)** Effect of α-synuclein concentration on the condensate partitioning of synapsin. Note the higher partitioning values in comparison to the measurements of α-synuclein partitioning (Figure 5F)

**Figure S10**

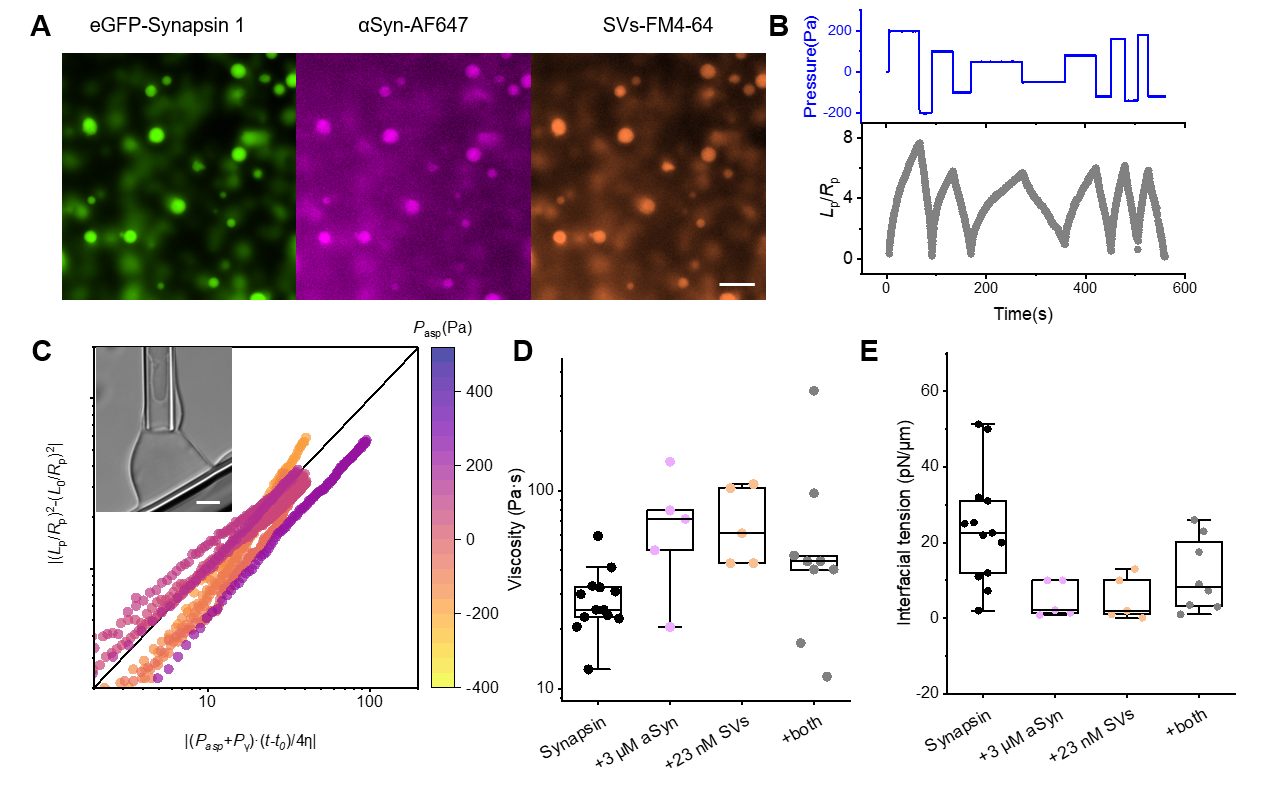

**Figure S10. MPA of synapsin/SV/α-synuclein condensates.**

**(A)** Fluorescence images showing the colocalization of eGFP-synapsin 1, α-synuclein-AF647, and SVs-FM 4-64. Scale bar: 5 μm. **(B)** Aspiration pressure (upper) and normalized aspiration length (lower) during MPA of condensates reconstituted in 9 μM synapsin, 3 μM α-synuclein and 23 nM SVs. **(C)** The plot of rescaled and segmented data according to ref^1^. The color of the traces represents applied aspiration pressure. The x axis is rescaled time, and the y axis is shifted (*L*_p_/*R*_p_)^2^. The black lines are y = x. The transmitted light image shows a micropipette-aspirated synapsin/α-synuclein/SV condensate. Scale bars, 5μm. The viscosity (**D)** and interfacial tension (**E)** of synapsin condensate in four conditions: synapsin only, synapsin + 3 μM α-synuclein, synapsin + 23 nM SVs, synapsin + 3 μM α-synuclein + 23 nM SVs (both). Two-way ANOVA test revealed a significant interaction between αSyn and SVs on the interfacial tension of condensates (*p* = 0.005).

**Movie S1. MAPAC of the condensate in Figure 1D.**

**Movie S2. MAPAC of the condensate in Figure 2C.**

**Table S1: Size distribution of cytosolic biomolecular condensates based on published images in the literature.**

Currently, high accuracy MAPAC quantifications cannot be achieved on condensates highlighted in gray (condensate diameter consistently smaller than 1.5 µm, with darker shades indicating more challenging condensates for MAPAC).

| **Condensate**  **forming protein** | **Cell Type** | **d_max_ (µm)** | **d_avg_ (µm)** | **S.D.** | **n** | **Reference** |
| --- | --- | --- | --- | --- | --- | --- |
| P granules | C. elegans germ cells | **4** | 2.95 | 0.8 | 3 | Brangwynne et al (2009) |
| optoDDX4 | HEK293T | **3** | 1.7 | 0.65 | 10 | Shin et al (2018) |
| optoFUS | HEK293T | **3** | 1.42 | 0.6 | 10 | Shin et al (2018) |
| LAF-1 | C. elegans embryos | **1.4** | 0.94 | 0.4 | 7 | Elbaum-Garfinkle et al (2015) |
| RGG-GFP-RGG | HEK293T | **5** | 3.54 | 0.86 | 6 | Schuster et al (2018) |
| Stress granules | HEK293T | **12** | 8 | 2 | 7 | Fang et al (2019) |
| Stress granules  (G3BP1) | U2OS | **2.8 5** | 2 3.4 | 0.4 1 | 9 6 | Wheeler et al (2016) Jalihal et al (2020) |
| TDP-43 | HEK293XT | **3.1** | 2 | 0.6 | 5 | Fang et al (2019) |
| TDP-43 | CV-B NPC | **3.1** | 2.5 | 0.4 | 3 | Fang et al (2019) |
| TDP-43 | N2a | **4.35** | 2.54 | 0.76 | 11 | Watanabe et al. (2020) |
| TDP-43PrLD | HeLa | **4.34** | 2.82 | 1.08 | 6 | Dhakal et al. (2023) |
| HNRNPA2B1 | HEK293XT | **2.3** | 2 | 0.3 | 7 | Fang et al (2019) |
| HNRNPA2B1 | CV-B NPC | **1.7** | 1.4 | 0.2 | 9 | Fang et al (2019) |
| DCP1A | U2OS | **5** | 3.8 | 1 | 9 | Jalihal et al (2020) |
| Balbiani body | Oocytes | **50** | 50 | N.A. | 1 | Boke et al (2016) |
| SPD-5 | C. elegans embryos | **2.5** | 2.5 | N.A. | 2 | Woodruff et al (2017) |
| ZO1 | MDCK | **9** | 6.7 | 1.5 | 4 | Beutel et al (2019) |
| ZO2 | MDCK | **4.5** | 2.5 | 1 | 6 | Beutel et al (2019) |
| ZO3 | MDCK | **1.4** | 0.8 | 0.4 | 16 | Beutel et al (2019) |
| NPR1 | Col-0 seedling | **3.1** | 2.1 | 0.6 | 11 | Zavaliev et al (2020) |
| sim3 | Col-0 seedling | **4** | 2.9 | 1 | 12 | Zavaliev et al (2020) |
| FUS (normal) | HeLa | **1.1** | 0.7 | 0.2 | 5 | Patel et al (2015) |
| FUS (stress) | HeLa | **2.1** | 1.4 | 0.6 | 12 | Patel et al (2015) |
| 3xFUS | HEK 293T | **1.78** | 1.25 | 0.31 | 16 | Brumbaugh-Reed et al. (2024) |
| FUS CHOP oncoprotein | NIH3T3 | **4.14** | 2.69 | 0.45 | 44 | Brumbaugh-Reed et al. (2024) |
| DDx4 | HeLa | **2.7** | 2.1 | 0.3 | 7 | Nott et al (2015) |
| whi3 | Ashbya | **1** | 0.65 | 0.2 | 7 | Zhang et al (2015) |
| Tau | Neuron | **3.8** | 2.4 | 0.8 | 4 | Wegmann et al (2018) |
| Huntingtin exon1 | HEK 293T | **5.3** | 2.3 | 1.2 | 4 | Peskett (2018) |
| α-synuclein | HeLa | **0.81** | 0.61 | 0.08 | 940 | Ray et al (2020) |
| α-Synuclein-5Fm | HeLa, SH-SY5Y | **10.48** | 2.99 | 1.9 | 51 | Prioska et al. (2023) |
| Synapsin +α-synuclein | Neuron HEK293T | **2.2 4.2** | 1.5 2.2 | 0.2 1.3 | 6 19 | Hoffmann et al (2021) |
| Synapsinα+α-synuclein | HEK293T | **4.8** | 1.9 | 1 | 75 | Current study |
| TFG | HeLa | **6.06** | 2.1 | 1.7 | 7 | Wegeng et al. (2024) |
| PSD-95 | HeLa | **2.8** | 1.8 | 0.6 | 5 | Zhu et al. (2024) |
| GlyR-βLD | HeLa | **2.6** | 2.6 | N.A. | 1 | Zhu et al. (2024) |
| TIAR-2 | PLM axon | **1.81** | 1 | 0.5 | 4 | Andrusiak et al. (2019) |
| M7ANC/USH1C/USH1G | HeLa | **2.29** | 1.3 | 0.6 | 8 | He et al. (2019) |
| M7BNC/USH1C/ANKS4B | HeLa | **2.62** | 1.26 | 0.97 | 6 | He et al. (2019) |
| Q74 (aggregates) | HEK293 | **15** | 8.58 | 3.4 | 7 | Llamas et al. (2023) |
| NEMO | U2OS | **1.9** | 1.2 | 0.4 | 7 | Du et al. (2022) |
| GPSM2 | HEK293 | **4.7** | 2.3 | 1.6 | 6 | Shi et al. (2022) |
| EML4–ALK | HeLa | **1.1** | 0.89 | 0.22 | 2 | Qin et al. (2021) |
| YAP-M | C2C12 | **4.2** | 2.9 | 0.9 | 9 | Hu et al. (2023) |
| YAP-C | C2C12 | **3.1** | 2.4 | 0.5 | 3 | Hu et al. (2023) |
| CCNT1 | HeLa | **3.75** | 2.3 | 0.6 | 13 | Guo et al. (2020) |
| AFF4 | HeLa | **2.22** | 1.4 | 0.5 | 10 | Guo et al. (2020) |
| AFF4+CCNT1 | HeLa | **1.67** | 1.1 | 0.4 | 9 | Guo et al. (2020) |
| SPOP+DAXX | HeLa | **3** | 2.3 | 0.4 | 5 | Bouchard et al. (2018) |
| MxA | Huh7 | **4.5** | 2 | 1.14 | 10 | Davis et al. (2019) |
| NLRP6-dsRNA | MEF | **1.67** | 1.67 | N.A. | 1 | Shen et al. (2021) |
| LAT | Jurkat T | **0.9** | 0.9 | N.A. | 1 | Su et al. (2016) |
| Epac1 | HEK 293T | **3.48** | 0.94 | 0.47 | 107 | Yang et al. (2022) |
| Epac1-R279E | HEK 293T | **2.42** | 1.39 | 0.56 | 14 | Yang et al. (2022) |
| PRC1.6 | HEK 293T, U2OS, HOS, 143B | **5.96** | 2.83 | 1.43 | 31 | Zhong et al. (2023) |
| N-Myc | SH-EP | **0.98** | 0.69 | 0.13 | 46 | Yang et al.（2024） |

*d*_max_*: diameter of largest condensates, d*_avg_*: average condensate diameter; S.D.: standard deviations. n: number of condensates. N.A.: not applicable.*

**Supplementary Discussions**

**Viscoelastic models**

In the quest for a simple viscoelastic model that fits all MAPAC data on cellular synapsin condensates (Figure 2), we surveyed the most commonly used models for viscoelastic soft materials ^2-7^. Table S1 summarizes the constitutive equations and creep responses of these models.

The responses of the majority (53/93) of cellular synapsin/α-synuclein condensates resemble the red trace in Figure 2A and have the following two key features: 1, a fast nonlinear increase in the aspiration length (*L*_p_) upon pressure application; 2, continued slow increase in *L*_p_ under sustained pressure, resulting in a permanent deformation after pressure removal. 24/93 condensates violated feature 1, where *L*_p_ increased linearly throughout the applied pressure (Figure 2A, black). 16/93 condensates violated feature 2, where neither long-term flow nor permanent deformation were observed (Figure 2A, blue). We aim to find the simplest model that captures the two key features while allowing apparent violations to these features due to the limitation of experimental resolutions.

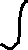

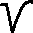

Feature 1 calls for the inclusion of an elastic component while feature 2 requires the long-term response of the model to be liquid. Therefore, the simplest model would appear to be the 2-element Maxwell model (Table S1), where an elastic spring is in series with a viscous dashpot. However, the Maxwell model does not allow materials to be purely viscous (Figure 2A, black): Setting *E* = 0 breaks the model and does not allow a dashpot with nonzero viscosity to respond. Furthermore, apparent viscous materials with minimal elastic response ($\frac{\sigma_{0}}{E}\to0$) would have a near infinite elasticity ($E\to\infty$ ). Additionally, the short-term nonlinear responses we observed (Figure 2A, red) are typically not abrupt and slower than the pressure change. For these reasons, the 2-element Maxwell model is not sufficient to explain our data.

Next, we evaluate 3-element viscoelastic models. There are two variations (the Maxwell-form and the Kelvin-form) for both the 3-element solid and the 3-element liquid models (Table 1). It has been shown that the two forms of the solid model are mathematically equivalent ^6^. Following terminology in Table 1 and using the superscript “M” and “K” to represent the components in the Maxwell-from and the Kelvin-form, respectively, it can be shown that setting $E_{1}^{M}=\frac{E_{1}^{K}E_{2}^{K}}{E_{1}^{K}+E_{2}^{K}}$ , $E_{2}^{M}=\frac{\left( E_{1}^{K} \right)^{2}}{E_{1}^{K}+E_{2}^{K}}$ , and $\eta^{M}=\frac{\eta^{K}\cdot\left( E_{1}^{K} \right)^{2}}{\left( E_{1}^{K}+E_{2}^{K} \right)^{2}}$ converts the constitutive equation for the Maxwell-form ($\sigma+\frac{\eta^{M}}{E_{2}^{M}}\dot{\sigma}=E_{1}^{M}\varepsilon+\frac{\eta(E_{1}^{M}+E_{2}^{M})}{E_{2}^{M}}\dot{\varepsilon}$) to that of the Kelvin form ($\sigma+\frac{\eta^{K}}{E_{1}^{K}+E_{2}^{K}}\dot{\sigma}=\frac{E_{1}^{K}E_{2}^{K}}{E_{1}^{K}+E_{2}^{K}}\varepsilon+\frac{E_{1}^{K}\eta^{K}}{E_{1}^{K}+E_{2}^{K}}\dot{\varepsilon}$) ^6^. Similarly, for the 3-element liquid models, setting $\eta_{1}^{M}=\frac{\eta_{1}^{K}\cdot\eta_{2}^{K}}{\eta_{1}^{K}+\eta_{2}^{K}}$, $\eta_{2}^{M}=\frac{\left( \eta_{1}^{K} \right)^{2}}{\eta_{1}^{K}+\eta_{2}^{K}}$, and $E^{M}=\left( \frac{\eta_{1}^{K}}{\eta_{1}^{K}+\eta_{2}^{K}} \right)^{2}E^{K}$ will convert the Maxwell-form to the Kelvin-form.

Feature 2 suggests a greater applicability of the 3-element liquid models (also known as Jeffreys Models ^5^) to our data, as these models include an independent dashpot that allows permanent deformations after stress removal. Setting *E* = 0 in a 3-element liquid model leads to a purely viscous material that resembles the black trace in Figure 2A. On the other hand, condensates that appear solid (Figure 2A, blue) can be treated as a liquid with near-infinite viscosity (a liquid that did not flow beyond one pixel under the pressure and timeframe of the experiment). Therefore, 3-element liquid models are sufficient to explain our data. We chose the Kelvin-form because individual elements in this model have more direct physical meanings as compared to in the Maxwell-form ^5, 6^.

Our current data cannot distinguish a 3-element liquid model from more complex models (e.g., 4-element models) ^2^. Therefore, we did not choose 4-element or higher-order models to avoid overfitting.

| **Name** | **Constitutive equation** | **Spring-dashpot representations** | **Strain response to a stepwise stress (blue)**  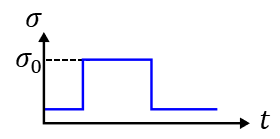 |
| --- | --- | --- | --- |
| Maxwell model | $\sigma+\frac{\eta}{E}\dot{\sigma}=\eta\dot{\varepsilon}$ | 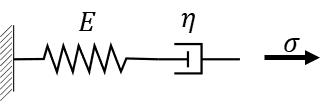 | 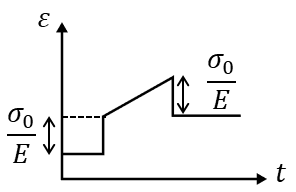 |
| Kelvin -Voigt Model | $\sigma=E\eta+\eta\dot{\varepsilon}$ | 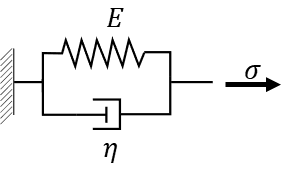 | 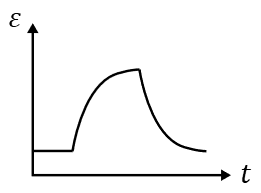 |
| **Three-element liquid -Kelvin form** | $\sigma+\frac{\eta_{1}+\eta_{2}}{E}\dot{\sigma}=\eta_{1}\dot{\varepsilon}+\frac{\eta_{1}\eta_{2}}{E}\ddot{\varepsilon}$ | 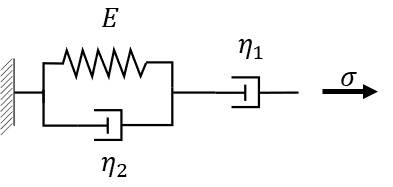 | 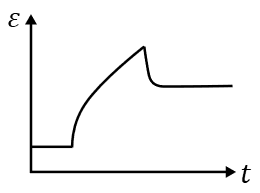 |
| Three-element liquid -Maxwell form | $\sigma+\frac{\eta_{2}}{E}\dot{\sigma}={(\eta}_{1}+\eta_{2})\dot{\varepsilon}+\frac{\eta_{1}\eta_{2}}{E}\ddot{\varepsilon}$ | 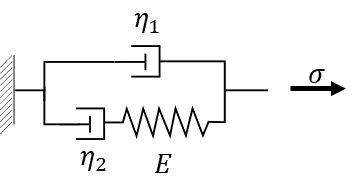 |  |
| Three-element solid -Kelvin form | $\sigma+\frac{\eta}{E_{1}+E_{2}}\dot{\sigma}=\frac{E_{1}E_{2}}{E_{1}+E_{2}}\varepsilon+\frac{E_{1}\eta}{E_{1}+E_{2}}\dot{\varepsilon}$ | 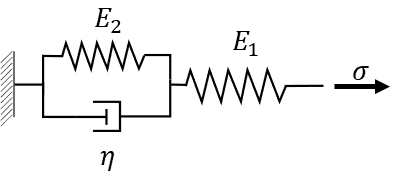 | 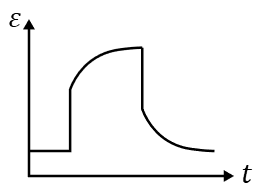 |
| Three-element solid -Maxwell form | $\sigma+\frac{\eta}{E_{2}}\dot{\sigma}=E_{1}\varepsilon+\frac{\eta(E_{1}+E_{2})}{E_{2}}\dot{\varepsilon}$ | 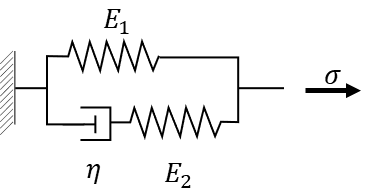 |  |
| Four element model | $\sigma+\left( \frac{\eta_{1}}{E_{1}}+\frac{\eta_{2}}{E_{2}}+\frac{\eta_{2}}{E_{1}} \right)\dot{\sigma}+\frac{\eta_{2}E_{1}}{E_{2}\eta_{1}}\ddot{\sigma}=\eta_{1}\dot{\varepsilon}+(\frac{\eta_{1}\eta_{2}}{E_{1}}+\frac{\eta_{1}\eta_{2}}{E_{2}})\ddot{\varepsilon}$ | 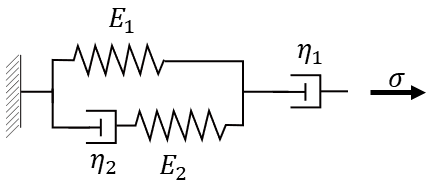 | 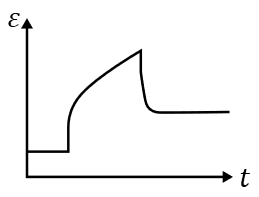 |

**Table S1: Linear viscoelastic models**

**Derivation and solution of the constitutive equation for 3-element liquid model-Kelvin form**

For the Kelvin form of the 3-element liquid model (Table S1, Figure 2B), the independent dashpot has a viscosity of $\eta_{1}$, the spring has an elasticity of $E$, and the dashpot in the Kelvin part has a viscosity of $\eta_{2}$ *.* Stress ($\sigma$) and strain ($\varepsilon$) on individual components are described by: $\sigma_{D_{1}}=\eta_{1}\dot{\varepsilon_{D_{1}}}$, $\sigma_{S}=E\varepsilon_{S}$, and $\sigma_{D_{2}}=\eta_{2}\dot{\varepsilon_{D_{2}}}$. The subscript “D_1_” represents the dashpot that has a viscosity of $\eta_{1}$, “S” represents the spring, “D_2_” represents the dashpot that has a viscosity of $\eta_{2}$.

Because the Kelvin part and the dashpot are in series, we have,

$$\sigma=\sigma_{K}=\sigma_{D_{1}} (S1)$$

$$\varepsilon=\varepsilon_{K}+\varepsilon_{D_{1}} (S2)$$

The subscript “K” represents the Kelvin part of the three-element model.

In the Kelvin part, we have,

$$\sigma_{K}=\sigma_{S}+\sigma_{D_{2}} \left( S3 \right)$$

$$\varepsilon_{K}=\varepsilon_{D_{2}}=\varepsilon_{S} (S4)$$

From eqs. S1, S3, and S4, we have,

$${\sigma=\sigma}_{K}=E\varepsilon_{K}+\eta_{2}\dot{\varepsilon_{K}} \left( S5 \right)$$

Differentiate eq. S2, and use $\dot{\varepsilon_{D_{1}}}=\frac{\sigma_{D_{1}}}{\eta_{1}}=\frac{\sigma}{\eta_{1}}$, we have,

$$\dot{\varepsilon_{K}}=\dot{\varepsilon}-\dot{\varepsilon_{D_{1}}}=\dot{\varepsilon}-\frac{\sigma}{\eta_{1}} (S6)$$

Combine eq. S2, S5, and S6,

$$\sigma=E\left( \varepsilon-\varepsilon_{D_{1}} \right)+\eta_{2}\left( \dot{\varepsilon}-\frac{\sigma}{\eta_{1}} \right) \left( S7.1 \right)$$

Differentiate eq. S7.1, and use $\dot{\varepsilon_{D_{1}}}=\frac{\sigma}{\eta_{1}}$,

$$\dot{\sigma}=E\left( \dot{\varepsilon}-\frac{\sigma}{\eta_{1}} \right)+\eta_{2}\left( \ddot{\varepsilon}-\frac{\dot{\sigma}}{\eta_{1}} \right) \left( S7.2 \right)$$

Rearrange eq. S7.2, we get the constitutive equation for the Kelvin form of the 3-element liquid model (Table S1):

$$\sigma+\frac{\eta_{1}+\eta_{2}}{E}\dot{\sigma}=\eta_{1}\dot{\varepsilon}+\frac{\eta_{1}\eta_{2}}{E}\ddot{\varepsilon} (S7.3)$$

Under a constant stress (resembling each pressure step in our experiments, Figure 2): $\sigma=\sigma_{0}$, $\dot{\sigma}=0$

Equation S7.3 becomes,

$$\sigma_{0}=\eta_{1}\dot{\varepsilon}+\frac{\eta_{1}\eta_{2}}{E}\ddot{\varepsilon} (S8)$$

Integrate eq. S8,

$$\sigma_{0}t+C_{1}=\eta_{1}\varepsilon+\frac{\eta_{1}\eta_{2}}{E}\dot{\varepsilon} (S9)$$

$C_{1}$ is a constant.

Divide both sides by $\frac{\eta_{1}\eta_{2}}{E}$,

$$\frac{\sigma_{0}E}{\eta_{1}\eta_{2}}t+\frac{EC_{1}}{\eta_{1}\eta_{2}}=\dot{\varepsilon} +\frac{E}{\eta_{2}}\varepsilon(S10)$$

Multiple eq. S10 with the integration factor $M\left( t \right)=e^{\int\frac{E}{\eta_{2}}dt}=e^{\frac{E}{\eta_{2}}t}$,

$$e^{\frac{E}{\eta_{2}}t}\cdot\frac{\sigma_{0}E}{\eta_{1}\eta_{2}}t+e^{\frac{E}{\eta_{2}}t}\cdot\frac{EC_{1}}{\eta_{1}\eta_{2}}=e^{\frac{E}{\eta_{2}}t}\cdot\dot{\varepsilon} +\varepsilon\cdot\frac{E}{\eta_{2}}e^{\frac{E}{\eta_{2}}t}=\frac{d\left( e^{\frac{E}{\eta_{2}}t}\varepsilon\right)}{dt} (S11)$$

Integrate eq. S11,

$$e^{\frac{E}{\eta_{2}}t}\varepsilon=e^{\frac{E}{\eta_{2}}t}\cdot\left( \frac{\sigma_{0}}{\eta_{1}}t-\frac{\sigma_{0}\eta_{2}}{\eta_{1}E} \right)+e^{\frac{E}{\eta_{2}}t}\cdot\frac{C_{1}}{\eta_{1}}+C_{2} (S12)$$

$C_{2}$ is a constant.

Divide both sides of eq. S12 by $e^{\frac{E}{\eta_{2}}t}$,

$$\varepsilon= \frac{\sigma_{0}}{\eta_{1}}t-\frac{\sigma_{0}\eta_{2}}{\eta_{1}E}+\frac{C_{1}}{\eta_{1}}+C_{2}e^{-\frac{E}{\eta_{2}}t} (S13)$$

When t = 0, $\varepsilon\left( 0 \right)=0,$

$$-\frac{\sigma_{0}\eta_{2}}{\eta_{1}E}+\frac{C_{1}}{\eta_{1}}+C_{2}=0 (S14)$$

When t is large, $\varepsilon\left( t\to\infty\right)=\frac{\sigma_{0}}{\eta_{1}}t+\frac{\sigma_{0}}{E}$,

$$-\frac{\sigma_{0}\eta_{2}}{\eta_{1}E}+\frac{C_{1}}{\eta_{1}}=\frac{\sigma_{0}}{E} (S15)$$

Combine eq. S13 to S15, we have:

$$\varepsilon= \frac{\sigma_{0}}{\eta_{1}}t+\frac{\sigma_{0}}{E} \left( 1-e^{-\frac{E}{\eta_{2}}t} \right) (S16.1)$$

If we use a more general initial condition: at $t= t_{0}$, $\varepsilon\left( 0 \right)=\varepsilon_{0},$

$$\varepsilon=\varepsilon_{0}+ \frac{\sigma_{0}}{\eta_{1}}\left( t-t_{0} \right)+\frac{\sigma_{0}}{E} \left( 1-e^{-\frac{E}{\eta_{2}}\left( t-t_{0} \right)} \right) (S16.2)$$

**Fitting MAPAC data to the 3-element liquid viscoelastic model**

Equation S16 describes the time dependent strain ($\varepsilon$) under a constant stress ($\sigma_{0}$) according to the Kelvin form of the 3-element liquid model. Next, we need to correlation stress and strain in the model to the pressure and condensate deformation in our MAPAC measurements.

The stress in our system is the aspiration pressure that induces condensate flow, therefore,

$$\sigma_{0}=P_{\mathrm{asp}}-P_{\gamma} (S17)$$

Where $P_{\gamma}=2\gamma\left( \frac{1}{R_{p}}-\frac{1}{R_{c}} \right)$ is the capillary pressure that needs to be overcome to initiate condensate flow. *R*_p_ and *R*_c_ are the radii of the micropipette and the condensates, respectively (Figure 1A).

We define strain as the normalized aspiration length (*L*_p_; Figure 1A):

$$\varepsilon=\frac{L_{p}}{R_{p}} (S18)$$

Therefore, eq. S16.2 becomes,

$$\frac{L_{p}}{R_{p}}=\left( \frac{L_{p}}{R_{p}} \right)_{0}+\frac{\left( P_{\mathrm{asp}}-P_{\gamma} \right)}{E}\left[ 1-e^{-\frac{E}{\eta_{2}}\left( t-t_{0} \right)} \right]+\frac{\left( P_{\mathrm{aps}}-P_{\gamma} \right)}{\eta_{1}}\left( t-t_{0} \right)$$

$$= \left( \frac{L_{p}}{R_{p}} \right)_{0}+A\left[ 1-e^{-\frac{(t-t_{0})}{\tau}} \right]+S\left( t-t_{0} \right) (S19)$$

Which is the fitting equation (eq. 1) described in the main text. This equation has three fitting parameters: $A=\frac{P_{\mathrm{asp}}-P_{\gamma}}{E}$, $B=\frac{P_{\mathrm{asp}}-P_{\gamma}}{\eta_{1}}$, and $\tau=\frac{\eta_{2}}{E}$. By measuring the response of *L*_p_ under different pressure, we can get relations between “*P*_asp_ vs. *B*” and “*P*_asp_ vs. *A*” for the condensate (Figure 2D, 2E). The slope of “*P*_asp_ vs. *B*” gives *η*_1_, while the slope of “*P*_asp_ vs. *A*” gives *E*. The intercept of the two relations gives *P*_γ_.

While *P*_γ_ can be converted to the interfacial tension of condensates following $P_{\gamma}=2\gamma\left( \frac{1}{R_{p}}-\frac{1}{R_{c}} \right)$. The relation between *η*_1_, *E*, and the viscoelasticity of condensates requires additional considerations. Here we follow a similar approach as in Guevorkian et al ^2^.

Cellular synapsin condensates do not wet the inner wall of the micropipette (Figure S2). Therefore, the recently developed calibration-free model^1^ for in vitro condensates that wet the micropipette cannot be directly applied here. This is further supported by a near linear long-term response of *L*_p_ under constant pressure (Figure 1E, 2A), in contrast to the *L*_p_ ~ t^0.5^ relationship predicted for condensates that wet the micropipette^1^. Following simulations of micropipette-aspirated cells with slipping boundary conditions^2, 8^, we used a linear conversion factor of 6 between *η*_1_ and condensate viscosity^4, 8^ and a linear conversion factor of 1 between *E* and the elasticity of condensates^2, 9^:

$$\mathrm{Viscosity}=\frac{\eta_{1}}{6}$$

$$\mathrm{Elasticity}=E$$

It is worth emphasizing that many simplifications were used in deriving the conversion factors between viscoelasticity and *η*_1_, *E*. The conversion factor between viscosity and $\eta_{1}$ range from 2 to 3π, depending on the choice of models ^2, 4, 8, 10^. Therefore, the viscosity of cellular condensates (Figure 2) may be off by up to 3-fold when using 6 as the conversion factor. Further theoretical and simulation efforts are needed to better understand the hydrodynamics of non-wetting viscoelastic condensates in a micropipette. Finally, cellular condensates can be heterogeneous or undergo shear thickening/thinning, these factors may also contribute to inaccuracies in our measurements of cellular condensates.

When fitting eq. S19 to cellular condensates, large fitting uncertainties were observed for data where the signal-to-noise ratio was low, the pressure profile significantly deviated from a step-function, or the pressure step was not sufficiently long (Figure S3). Following Guevorkian et al ^2^, we evaluate a linearized version of eq. S19 with only 2 fitting parameters, focusing on the long-term response of *L*_p_ (Figure 2B, 2C). If we assume *E* is not negligible, the long-term response of eq. S19 follows a linear relation:

$$\frac{L_{p}}{R_{p}}=\left( \frac{L_{p}}{R_{p}} \right)_{0}+A+B\left( t-t_{0} \right) (S20.1)$$

with only 2 fitting parameters: $A=\frac{P_{\mathrm{aps}}-P_{\gamma}}{E}$, $B=\frac{P_{\mathrm{aps}}-P_{\gamma}}{\eta_{1}}$.

The resulting “*P*_asp_ vs. *B*” and “*P*_asp_ vs. *A*” relations from the linear 2-parameter model were indistinguishable from those of the full 3-parameter model (Figure 2D, 2E, Figure S3). However, by sacrificing the fitting for *η*_2_ (Figure S3F), a property not directly related to the material state of the condensate ^5, 6^, the linear model is more stable and reduced fitting errors.

For condensates that have an apparent elastic response but no measurable flow (Figure 2A, blue), we assumed that the applied pressure step was not long enough to induce a detectable amount of flow (1 pixel). Assuming *T*_p_ is the duration of a constant aspiration pressure *P*_max_ (*P*_max_>> *P*_γ_) and the pixel length is Δ, we assign $\mathrm{Viscosity}=\frac{P_{\max}T_{p}R_{p}}{6\Delta}$ for those condensates. If the elastic response of the condensate is also below 1 pixel (Figure 3D, cell #2), we assign $\mathrm{Elasticity}=\frac{P_{\max}R_{p}}{\Delta}$.

For condensates that did not show measurable elastic deformation but showed clear flow (e.g., Figure 2A, black), there are two scenarios:

#1) The condensate has near infinite elasticity ($E\to\infty$), but finite viscosity. eq. S19 becomes,

$$\frac{L_{p}}{R_{p}}=\left( \frac{L_{p}}{R_{p}} \right)_{0}+\frac{\left( P_{\mathrm{asp}}-P_{\gamma} \right)}{\eta_{1}}\left( t-t_{0} \right)$$

$$=\left( \frac{L_{p}}{R_{p}} \right)_{0}+B\left( t-t_{0} \right) (S20.2)$$

With 1 fitting parameter $B=\frac{P_{\mathrm{asp}}-P_{\gamma}}{\eta_{1}}$. Here, “*P*_asp_ vs. *B*” again gives $\eta_{1}$ and the viscosity of the condensate is $\frac{\eta_{1}}{6}$. Following previous discussions, we assign $\mathrm{Elasticity}=\frac{P_{\max}R_{p}}{\Delta}$ for these condensates.

#2) The condensate has negligible elasticity ($E\to0$) but measurable viscosity. Here, the linearization of eq. 19 becomes,

$$\frac{L_{p}}{R_{p}}=\left( \frac{L_{p}}{R_{p}} \right)_{0}+\frac{\left( P_{\mathrm{asp}}-P_{\gamma} \right)}{\eta_{2}}\left( t-t_{0} \right) +\frac{\left( P_{\mathrm{asp}}-P_{\gamma} \right)}{\eta_{1}}\left( t-t_{0} \right)$$

$$=\left( \frac{L_{p}}{R_{p}} \right)_{0}+B_{\mathrm{obs}}\left( t-t_{0} \right) (S20.3)$$

with 1 fitting parameter: $B_{\mathrm{obs}}=\left( P_{\mathrm{asp}}-P_{\gamma} \right)\frac{1}{\eta_{\mathrm{obs}}}$, and $\frac{1}{\eta_{\mathrm{obs}}}=\left( \frac{1}{\eta_{1}}+\frac{1}{\eta_{2}} \right)$.

The observed viscosity ($\eta_{\mathrm{obs}}$) from the slope of “*P*_asp_ vs. *B*_obs_” is typically much smaller than $\eta_{2}$ (obtained from fitting high quality data to the full model, Figure S3F, Figure 3C). Therefore, $\mathrm{Viscosity}=\frac{\eta_{1}}{6}\approx\frac{\eta_{\mathrm{obs}}}{6}$ for these condensates. To assign condensate elasticity, we assume the maximal aspiration length (*L*_p_^max^ ~ 20 *R*_p_) did not cover the entire elastic response during the applied pressure step (*P*_min_ ~ 10 kPa). Therefore, we assign $\mathrm{Elasticity}=\frac{P_{\min}R_{p}}{{L_{p}}^{\max}}\sim500 \mathrm{Pa}$ for these condensates (Figure 2). The same argument applies to the evaluation of condensate elasticity in vitro (Figure 4 – Figure 6). In in vitro MPA measurements: *L*_p_^max^ ~ 10 *R*_p_ and *P*_min_ ~ 100 Pa. Therefore, the elasticity of in vitro condensates was assigned an elasticity of 10 Pa (Figure 5H).

Notably, the two scenarios can be loosely distinguished by whether the observed slope of “*P*_asp_ vs. *B*” is larger (scenario #1) or smaller (scenario #2) than the system viscosity $\eta_{2}$ obtained from fitting to the full model (Figure S3F, Figure 3C). All 24 synapsin/α-synuclein condensates that belonged to this category followed scenario #2 (Figure 2).

**Condensate partitioning**

When a client protein (e.g., α-synuclein) gets recruited into condensates (e.g., synapsin condensates), the binding isotherm follows:

$$c_{\mathrm{dense}}=\frac{c_{\max}}{1+\left( \frac{K_{D}}{c_{\mathrm{dillute}}} \right)^{N}} (S21)$$

Here, $c_{\mathrm{dense}}$is the concentration of the client in the condensate and $c_{\max}$ is the maximal concentration the client can reach in the condensate. $c_{\mathrm{dillute}}$ is the concentration of the client in the dilute phase. *K*_D_ is the dissociation constant, with *K*_D_^-1^ representing the affinity between the client and the dense phase. *N* represents homotropic cooperativity between client molecules, with *N* = 1 corresponding to independent binding between the client protein and the condensate.

The condensate partitioning coefficient of the client protein:

$$\Pi= \frac{c_{\mathrm{dense}}}{c_{\mathrm{dillute}}}=\frac{c_{\max}\cdot{c_{\mathrm{dillute}}}^{N-1}}{{c_{\mathrm{dillute}}}^{N}+{K_{D}}^{N}} (S22)$$

When, *N* = 1 (no cooperativity between client molecules)

$$\Pi= \frac{c_{\max}}{c_{\mathrm{dillute}}+K_{D}} (S23)$$

is a monotonic decreasing function of $c_{\mathrm{dillute}}$. However, $\Pi$ can increase with $c_{\mathrm{dillute}}$ if *N* is larger than 1. Therefore, the observed increase of $\Pi_{\alpha Syn}$ with α-synuclein concentration in the dilute phase (Figure 5F, S9C) indicates homotropic cooperativity (*N* > 1) between α-synuclein when getting recruited into synapsin condensates.

Notably, when $c_{\mathrm{dillute}}\ll K_{D}$, eq. S23 shows $\Pi\sim{K_{D}}^{-1}$, suggesting the condensate partitioning coefficient $\Pi$ can be directly used to approximate the affinity of the client protein towards the dense phase. More generally, $\log\left( \Pi\right)$ is linearly correlated with the free energy of condensate partitioning for the client protein ($-k_{B}T\log K_{D}$) via equation S24.

$$\log\left( \Pi\right)\approx\log c_{\max}\cdot{c_{\mathrm{dillute}}}^{N-1}-N\log K_{D} (S24)$$

On a technical note, unlike client concentrations estimated from fluorescence intensity measurements (Figure S9, S6A), the measurement of partitioning coefficient $\Pi$ is independent of heterogeneities in fluorescence excitation, or variations in intracellular environment (e.g. pH). Therefore, $\Pi$ is a robust parameter that can be used to infer the affinity of client molecules towards condensates.
